# Competition against the intestinal microbiota selects for pathoadaptive traits in ESBL *E. coli*

**DOI:** 10.64898/2026.06.25.730905

**Authors:** Pramod K. Jangir, Leonardo F. Lemos Rocha, Marco Molari, Nicolas Wenner, Cecilia Fruet, Pablo Manfredi, Carlos Flores, Anna Sintsova, Roxanne Mouchet, Lisa Diner, Anouk Bertola, Svenja Förster, Elisa Le, MiCayla Johnson, Malenka Kunz, Andrea Rocker, Adrian Egli, Urs Jenal, Christoph Dehio, Anne-Florence Bitbol, Médéric Diard

## Abstract

The intestinal tract is a reservoir for Extended-Spectrum β-Lactamase (ESBL)-producing *Escherichia coli*. Asymptomatic gut colonization by these pathobionts represents a major risk for extraintestinal infections. Despite clinical relevance, the genetic basis of gut colonization by ESBL *E. coli* remains poorly understood. Here, we determined how the microbiota shapes the fitness landscape of diverse ESBL *E. coli* strains, defining the functional requirements for intestinal colonization. In microbiota-depleted hosts, colonization relies mostly on metabolic functions. In contrast, in mice harbouring a microbiota, pathoadaptive functions associated with adhesion and biofilm formation are dominant determinants of *E. coli* fitness, together with accessory virulence functions. Consistent with these observations, experimental evolution in mice reveals convergent adaptation of ESBL *E. coli* to the presence of a complex microbiota through enhanced adhesion. These findings establish the microbiota as a major ecological driver of pathoadaptation in antibiotic-resistant pathobionts.

## Introduction

*Escherichia coli* is the leading cause of mortality associated with antimicrobial resistance worldwide^1,2^. Extended-spectrum β-lactamase (ESBL)-producing *E. coli* represent a particularly urgent public health concern^3–5^. A key feature of ESBL *E. coli* is their ability to asymptomatically colonize the intestinal tract, which serves as a reservoir for subsequent dissemination to the urinary tract or bloodstream^2,4,6,7^. However, how ESBL *E. coli* establish persistent gut colonization remains unclear.

The mammalian gut is a complex ecosystem in which the microbiota confers colonization resistance against invading pathogens through diverse mechanisms, including nutrient depletion, spatial exclusion, and the production of antimicrobials^8^. The prevailing working hypothesis is that successful gut colonization by ESBL *E. coli* is largely determined by metabolic capacity, whereby strains able to exploit available nutrients can achieve niche occupancy^9–12^. However, it was shown that metabolic flexibility alone may not explain the prevalence of certain ESBL *E. coli* clones^13^. The metabolic-centric view stems mainly from comparative genome analyses^10^, competition between closely related species or studies in experimental models lacking a complex gut microbial community^14,15^. Although the metabolic capacity is an important contributor to colonization, it is evident that the gut microbiota imposes diverse selective pressures on invading pathogens^8^ and that colonization success could depend on other traits.

Supporting this notion, insights from enteric pathogens provide evidence that fitness determinants in the gut are highly context-dependent and shaped by interactions with the gut microbiota, extending beyond nutrient competition. For example, in *Salmonella enterica*, virulence attenuated mutants expand in the microbiota-depleted gut, whereas virulent lineages are selected in the presence of the microbiota^16^. Similarly, virulence-associated traits are favoured in *Citrobacter rodentium* within hosts harbouring a competitive microbiota^17^. Together, these findings show that interactions with the gut microbiota can affect selection for diverse fitness and virulence-associated traits.

However, whether this microbiota-driven selection acts on pathobionts such as ESBL *E. coli*, which colonize the gut asymptomatically, and how it shapes colonization mechanisms, remains unclear. Notably, these strains are mainly Extra-intestinal Pathogenic *E. coli* (ExPEC) harbouring various virulence genes, many encoded on mobile genetic elements and pathogenicity islands, and previous studies have suggested that these genes could increase bacterial fitness during intestinal colonization^18^. Yet, it is unclear which functions of ESBL *E. coli* are required for successful colonization in the presence of a microbiota, and how microbiota-mediated competitions drive their evolution. Understanding these interactions is critical for designing effective prophylactic approaches against ESBL ExPEC, as the gut represents their primary ecological niche.

Here, we address these questions by combining genome-wide mutant fitness profiling with experimental evolution in mice across multiple pandemic ESBL *E. coli* lineages. By systematically comparing fitness determinants in microbiota-depleted and microbiota-colonized hosts, we reveal that the gut microbiota shapes the fitness landscape of ESBL *E. coli*, selectively favouring pathoadaptive traits, such as adhesion and biofilm formation, that are largely dispensable in microbiota-depleted hosts.

## Results

### The gut microbiota shifts the fitness landscape of ESBL *E. coli*

We first assessed how the gut microbiota shapes the fitness landscape of ESBL *E. coli*. For this, we performed comparative genome-wide fitness profiling in microbiota-colonized and microbiota-depleted mice using randomly barcoded transposon sequencing (RB-TnSeq). We generated RB-TnSeq libraries in four clinically relevant strains, isolated from healthy travelers (see Methods). These strains belong to globally prevalent sequence types (STs) representing pandemic multidrug-resistant *E. coli* clones^19^. We selected two ST131 strains from the subclones O25b FimH30 (EcST131 J) and O16 FimH41 (EcST131 I), one ST69 (EcST69), and one ST73 (EcST73) **(Figure 1A, Table S1)**. The libraries consisted of ∼326800 (EcST131 I), ∼53400 (EcST131 J), ∼27300 (EcST69), and ∼26100 (EcST73) unique single-insertion mutants, corresponding to ∼92%, ∼76%, ∼68%, and ∼63% coverage of annotated genomic features, respectively, including both genes and non-coding elements (e.g., rRNA, tRNA) **(Figure S1, Table S2)**. Across the strains, each feature was represented by 3-21 independent barcoded Tn*5* insertions, enabling robust fitness measurement **(Table S2)**.

**Figure 1.**
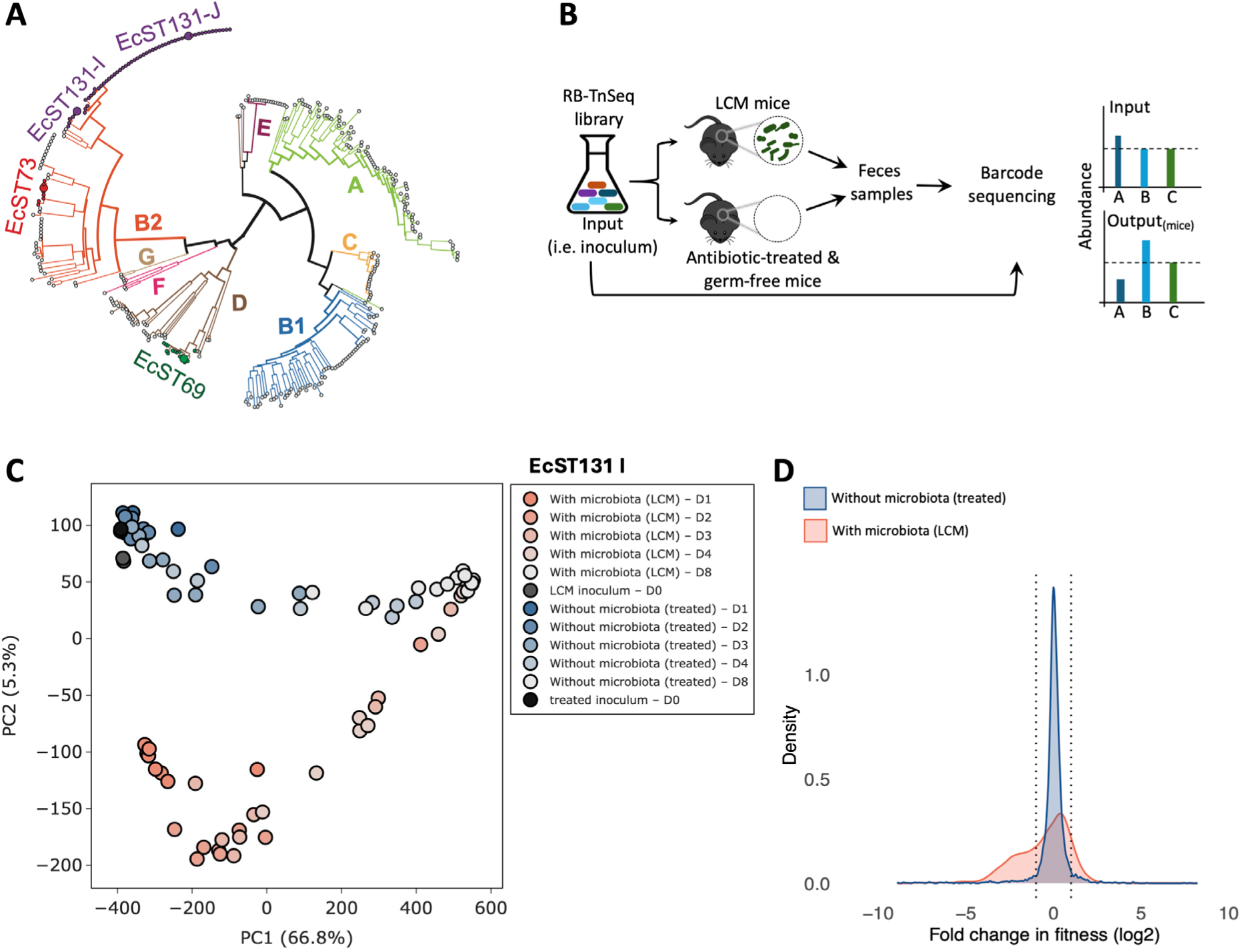
Gut microbiota alters *in vivo* fitness landscape of ESBL *E. coli*. **(A)** Phylogenetic placement of the four ESBL *E. coli* strains (highlighted) within a collection of 352 representative *E. coli* genomes spanning diverse STs (see the source data file). **(B)** Schematic overview of the RB-TnSeq experiments in mice with and without microbiota. **(C)** Principal component analysis of RB-TnSeq profiles of the strain EcST131 I across screening conditions, based on normalized transposon barcode abundances (see Methods). Each point represents an individual mouse sample. **(D)** Distribution of fitness effects (DFEs) of EcST131 I genes during bacterial colonization in the presence (red) and absence of the microbiota (blue; antibiotic-treated). Histograms are shown after Gaussian kernel density smoothing. Dashed lines indicate the neutral fitness window.

We performed comparative screening of these libraries using 25000-35000 mutants for each strain in low-complexity microbiota (LCM) and antibiotic-treated LCM mice (**Figure 1B**, see Methods). To identify effects of antibiotic treatment and to confirm that antibiotic-treated mice confidently recapitulate a microbiota-free niche, we additionally screened two of the strains (EcST131 J and EcST73) in germ-free (GF) mice. The LCM mice allow a stable and reproducible gut colonization, in which the presence of low-complexity microbiota enables large-scale screening of mutant libraries, while also revealing microbiota-dependent fitness effects^20,21^. Indeed, across all four strains, the overall bacterial load of the mutant libraries in LCM mice remained high (∼10^7^ CFU/g feces) and stable over time **(Figure S2)**. Importantly, the diversity of represented genomic features was well maintained during the early infection days (day 1 and/or day 2) but showed substantial loss at later time points **(Figure S3)**. The high similarity in diversity of genomic features between the input (before infection) and early output samples shows that the screen is feasible and not confounded by population bottlenecks or clonal expansion, as previously established to assess bottleneck effects in an *in vivo* transposon screen^22^. Therefore, we limited our RB-TnSeq analysis to day 1 or day 2 post-infection for a robust and reproducible dataset (see Methods). For each library, sequencing of two inocula shows high correlation (∼0.92 on average across strains and conditions), demonstrating reproducibility of library preparation, sequencing, and mapping (**Figure S4**).

Principal component analysis (PCA) highlighted consistent separation between pools of mutants from microbiota-colonized mice (LCM mice) and microbiota-depleted mice (antibiotic-treated and GF), indicating that the presence of the microbiota imposed distinct selective pressures on all four *E. coli* strains (**Figures 1C and S5A-C**). Of note, for the two strains also screened in GF mice, antibiotic-treated and GF mice samples clustered together, indicating that antibiotic-treated mice effectively recapitulate a microbiota-free environment (**Figures S5A and S5C**). Importantly, the clear separation of microbiota-colonized and microbiota-depleted mice samples on day 1 and 2 was followed by the convergence of the mutant pool compositions. This resulted from selective sweeps and reduced mutant diversity caused by the expansion of mutations conferring a strong fitness advantage across conditions (**Figure S3, Supplementary data file S1**).

The distribution of fitness effects (DFEs) of mutations differed between conditions (**Figures 1D and S5D-F**). In microbiota-colonized mice, DFEs were broader compared to antibiotic-treated and germ-free mice, indicating a greater number of both deleterious and beneficial mutations (**Figure S6**). In contrast, microbiota-depleted conditions (antibiotic-treated and GF) exhibited a narrow distribution centered near neutral fitness change, consistent with a less complex and more permissive environment (**Figures 1D, S5D-F, and S6**). These differences in the DFEs were consistent across all four strains, suggesting conserved responses to ecological constraints imposed by the microbiota (**Figure S5 and Supplementary data file S1**).

Altogether, these observations demonstrate that the microbiota substantially alters the fitness landscape of ESBL *E. coli* by increasing both the diversity and strength of selective pressures in the gut.

### The gut microbiota selects for pathoadaptive traits in ESBL *E. coli*

Next, we identified the genetic determinants of *E. coli* under selection by the presence of the microbiota. For this, we compared the fitness profiles of each transposon mutant between LCM mice and microbiota-depleted mice (antibiotic-treated LCM or GF). For more robust results, we restricted this analysis to intragenic insertions that directly disrupt genes, since the effects of intergenic insertions are uncertain, and it remains unclear which gene surrounding the insertion they primarily affect. Given that antibiotic-treated and GF samples clustered together (**Figure S5**), we considered both conditions as microbiota-depleted environments and included mutants showing fitness change in either condition. Pooling data from all four strains belonging to three prevalent STs enabled a robust genome-wide mutant fitness profiling, where signals observed with one strain could be confirmed or complemented by signals from the other strains. We found a total of 125 gene disruptions that reduced fitness in at least one strain, specifically when the microbiota was depleted (**Figure S7A**). This was substantially less than mutants showing reduced fitness exclusively in the presence of the microbiota (506 genes) (**Figure S8 and source data file**). We found 58 gene disruptions reducing fitness in both conditions (**Figure S7B**). This considerable difference between conditions indicated that the gut microbiota imposes a substantially broader and more specific set of functional requirements for colonization. Only genes for which a function could be assigned are shown in figures S7 and S8, meaning that a notable proportion (∼23%) of the mutations were in genes of unknown function **(Source data file)**.

In microbiota-depleted conditions, fitness was predominantly determined by metabolic functions (**Figures 2A and S7A**). Mutants defective in the uptake and utilization of readily available carbon sources, including mono- and di-saccharides arabinose (*araA and araB*), xylose (*xylA and xylB*), fructose (*fruA*), mannose (*manX and manZ*), melibiose (*melA*), sugar alcohol mannitol (*mtlD*) and hexuronates (*garK*, *kdgK*, and *uxuA*), showed significant fitness defects, suggesting sufficient luminal concentration of unabsorbed fermentable oligosaccharides, disaccharides, monosaccharides, and polyols (FODMAPs) in the absence of competitors. *E. coli* maintained redox balance and generated ATP anaerobically from these nutrients via the mixed-acid fermentation, evidenced by the requirement of the alcohol dehydrogenase (*adhE*) and phosphate acetyltransferase (*pta*); as well as via fumarate respiration (*frd* operon) (**Figure S7A**). In line with sustained growth to maintain high loads in this permissive environment, many genes involved in the biosynthesis and maintenance of nucleic acids and proteins were also required.

**Figure 2.**
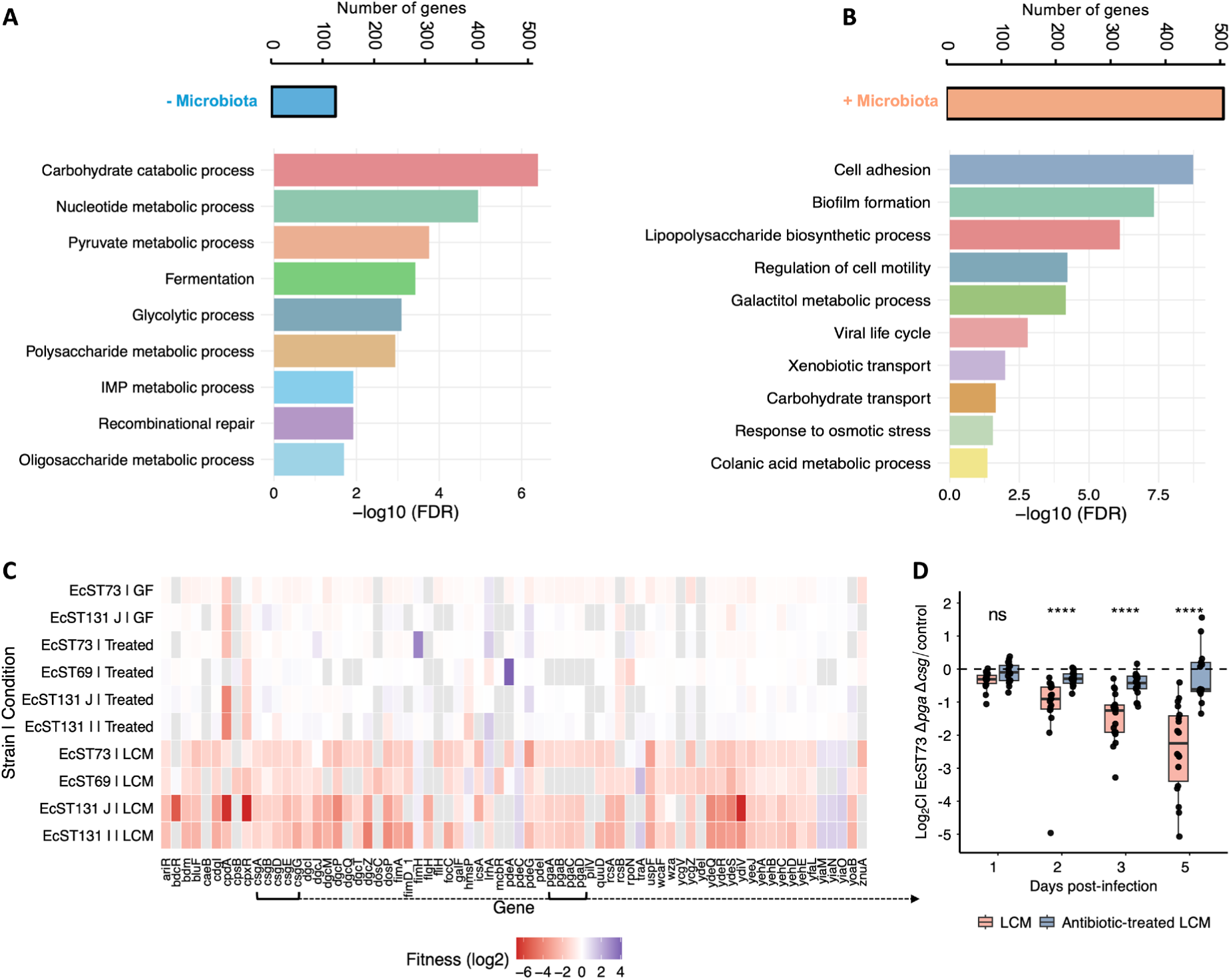
Biofilm and cell adhesion-associated pathoadaptive functions are specifically required for gut colonization in the presence of microbiota. Significantly, enriched “biological process” gene ontology terms among mutants showing fitness defect in the absence **(A)** and presence of microbiota **(B)**. The total number of genes in each category is shown in the bar plots at the top. *P* values were calculated using Fisher’s exact tests and multiple testing correction was performed using the Benjamini-Hochberg false discovery rate (FDR) method. **(C)** Heatmap showing RB-TnSeq-derived fitness scores for biofilm, cell adhesion, fimbriae and pili related genes that showed a significant fitness change in at least one strain in any condition screened (i.e., LCM, antibiotic-treated LCM and GF). **(D)** Competitive fitness of the EcST73 Δ*pga*Δ*csg* double mutant against the wild-type control. LCM and antibiotic-treated LCM mice were inoculated with 1:1 mixture of EcST73 wild-type and the mutant. Each data point represents one mouse, *n*=18 mice for LCM and *n*=17 for antibiotic-treated LCM from 3 independent experiments. Dotted line indicates the expected 1:1 ratio (equal fitness). *P* values were calculated using two-sided Wilcoxon rank-sum test and adjusted for multiple comparisons using the Holm method, \**p* <0.05, \*\**p* <0.001, \*\*\**p* <0.001, \*\*\*\**p* <0.0001.

Nevertheless, *E. coli* faced stress in the gut, such as bile acids, reflected by the requirement for core lipopolysaccharide (LPS) biosynthesis (*waa* genes) (**Figure S7B**). More specific to the absence of the microbiota, the counter-selection of mutations of the EnvZ/OmpR regulon, the CpxR system (envelope stress mitigation) and RpoS could result from the presence of osmotically active FODMAPs (**Figure S7A**). As expected, genes conferring resistance to ampicillin (such as *slt*^23^, *ampG*^24^, *dacA*^25^) were selectively required in antibiotic-treated but not in GF mice (**Figure S9**). Notably, despite differences in animal models and bacterial genetic background, many of genes identified here, including those involved in metabolism (such as *araA*, *araB*, *rbsK*, *malPQ*, *agaR*), respiration (*frdABD*) and stress resistance (such as *tolC*, *envZ*/*ompR*), have previously been linked to intestinal colonization in microbiota-depleted mice or rabbits, supporting the robustness of our screening approach^26–29^.

In contrast, the microbiota in LCM mice selected for distinct scavenging abilities and anaerobic respiration pathways, broader stress response, and importantly, pathoadaptive functions, including adhesion, biofilm formation, and virulence-associated traits in ESBL *E. coli* **(Figures 2B and S8).** The specific requirement of genes for the uptake and degradation of galactose (*mgl* genes), galactitol (*gat* operon), aryl-beta-glucosides (*bglH*), N,N’-diacetylchitobiose (GlcNAc2) (*chb* operon), glucuronides (*uidB and uidC*), and uncharacterized carbohydrates (*ydj* and *ycj* operons) suggested microbiota-mediated depletion of most carbon sources that *E. coli* preferentially used in its absence, and reliance on alternative ones, such as GlcNAc-containing sugars derived from mucus glycans^30^. The nutrient shift was confirmed by observing mutations that increased fitness in the presence of the microbiota (**Figure S7C**). For example, the utilization of xylose by *E. coli* was selected for in treated mice but was detrimental in LCM mice (**Figure S10A-B**), illustrating that optimization of carbon source utilization can be achieved by loss-of-function mutations. In this regard, the utilization of maltose and maltodextrins (*mal* genes) was strongly counter-selected during intestinal colonization with or without microbiota, as previously observed in mouse models^31–33^ and further validated here in competition experiments (**Figures S10C-F and S11**). Consistent with a resource-limited environment, genes involved in amino acid (*brnQ*, *tyrP*, and *ansP*) and peptide transport (*oppC*, *dtpA*, *mppA*, *dcp,* and *ptrB*), along with biosynthetic and catabolic enzymes, iron acquisition (*cirA*, *fepC*), as well as vitamin B12 uptake (*btuC*), were also required. For energy, *E. coli* relied on reductases for anaerobic respiration of nitrate (*nar* operons) and trimethylamine N-oxide (TMAO, *tor* operon) alongside requisite molybdopterin cofactor biosynthesis genes (**Figure S8**). As for *Salmonella enterica* Typhimurium during the initial colonization step in LCM mice^20^, di-hydrogen seemed to be a source of energy for *E. coli* in this context, with the selection for the hydrogenase 2 genes (*hybE, hybC, and hybO*).

The presence of the microbiota also imposed extra stresses, as evidenced by the requirement of multiple efflux pumps (*acrA*, *mdt,* and *emr*), pH homeostasis (*gadA/C*, *cadA*), osmotic stress tolerance (*osmB* and *osmC*), and membrane stability genes (e.g., *wbb* and *rfb* operon, *arnA* and *arnB*), probably to tolerate increased concentrations of competitors- and host-derived toxic metabolites and to resist sub-optimal acidic pH at high concentrations of short-chain fatty acids produced by the surrounding fermentative microbiota^34^.

Strikingly, genes involved in adhesion and biofilm formation were major determinants of fitness specifically in the presence of the microbiota (**Figures 2B and 2C**). Disruption of genetically distinct biofilm- and cell adhesion-associated pathways – including fimbriae (*fim* and *yeh*), curli fibers (*csg* operon), exopolysaccharide biosynthesis (*pgaABCD* genes and colonic acid biosynthesis *wca* genes), and c-di-GMP signaling (e.g., *dgcZ*, *dgcJ*) resulted in a significant fitness defect, indicating that the transition from a motile to sessile lifestyle, fine-tuned by c-di-GMP-mediated regulation, is critical under competitive conditions (**Figures 2C and S8**). To validate the role of biofilm- and cell adhesion-related genes in microbiota-colonized hosts, we constructed a double mutant lacking *pga* and *csg* operons in strain EcST73 (named EcST73 Δ*pga*Δ*csg*). The *pga* operon (involved in biosynthesis of poly-β-1,6-N-acetylglucosamine (PNAG)) and the *csg* operon (curli biosynthesis) play important roles in bacterial virulence^35,36^ and biofilm formation by distinct mechanisms. Specifically, PNAG forms an extracellular polysaccharide scaffold, whereas curli are amyloid fibers that mediate surface attachment and cell-cell interactions. Disrupting both systems, therefore, targets complementary components of biofilm formation, increasing the likelihood of robust phenotype *in vivo*. Consistent with this, competition experiments showed that the EcST73 Δ*pga*Δ*csg* mutant exhibited significantly reduced fitness in the presence of microbiota compared to the wild-type control (**Figure 2D**).

To determine whether the microbiota-specific fitness functions were associated with particular *E. coli* lineages, we analyzed the distribution of the 506 genes that increase fitness in the presence of microbiota across thousands of *E. coli* genomes, representing diverse STs and phylogroups. These genes were prevalent in strains from most STs belonging to phylogroup B2 (**Figure S12A**). Accessory genes were particularly represented (**Figures S13 and S14**), including multiple virulence factors, such as a hemolysin (*hlyE*), the cytolethal distending toxin (*cdt* operon), the colibactin (*clb* operon), and the siderophore Yersiniabactin (*irp1* and *irp2* genes) (**Figures S8, S12C, and S14**).

Together, these results demonstrate that ESBL *E. coli* relies on pathoadaptive functions, including accessory virulence traits, when competing with the microbiota during colonization of the mammalian gut. These functions extend beyond metabolic adaptation and include stress resistance, biofilm formation, adhesion and toxins.

### Evolution during prolonged gut colonization in the presence of a complex microbiota

To further understand ESBL *E. coli* adaptation to gut colonization in healthy hosts, that is, in the presence of a complex microbiota that drastically limits the effective population size of *E. coli*^37^, we shifted from LCM to Specific Pathogen Free (SPF) mice that harbour a conventional intestinal microbiota but lack detectable *Enterobacteriaceae* (**Figure S15**).

The four ESBL *E. coli* strains were able to colonize SPF mice for several months at reduced loads varying around 10^5^ CFU/g of feces (**Figure 3**), representing an approximately 100-fold reduction compared with LCM mice (**Figure S2A-D**). In contrast, the laboratory strain *E. coli* MG1655 failed to stably colonize SPF mice (**Figure S16**).

**Figure 3.**
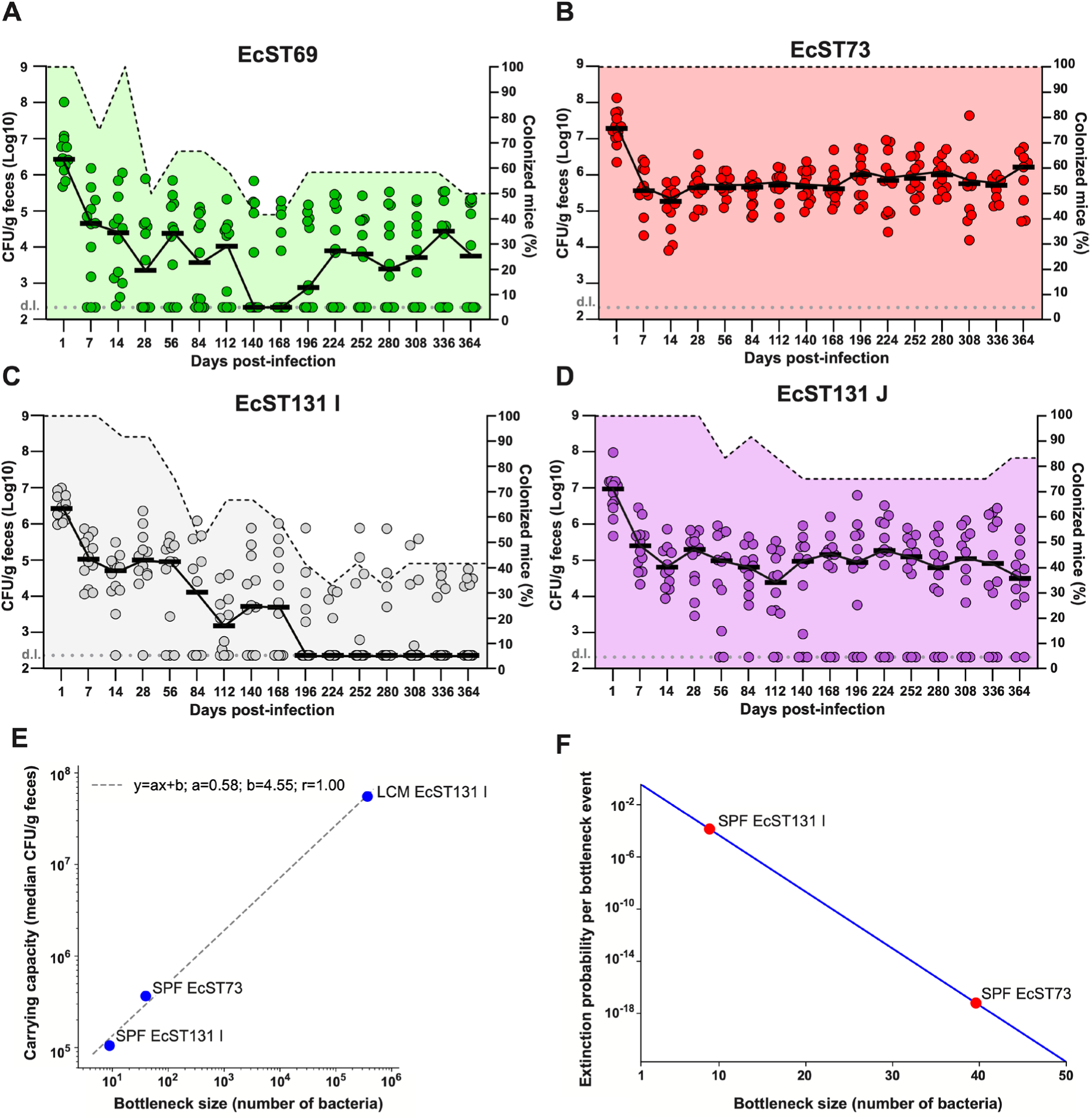
Long-term colonization dynamics of ESBL *E. coli* strains in conventional SPF mice. **(A-D)** Figures show fecal bacterial load for each ESBL *E. coli* during a 1-year *in vivo* evolution experiment in SPF mice harboring complex gut microbiota. Each point represents the bacterial load from an individual mouse at the indicated time points (*n*=12 mice per strain, from four independent experiments). Black horizontal line indicates the median bacterial load across mice at each time point. Shaded area shows the proportion of colonized mice at each time point (y-axis on the right). d.l. stands for detection limit. **(E)** The measured carrying capacity (median over mice, expressed in CFU/g) is plotted versus the bottleneck estimated from our maximum-likelihood approach, in the three cases considered in the **Figure S19** (markers). Logarithmic scales are employed on both axes. A linear fit is performed in these logarithmic scales (dashed gray line), revealing a non-linear scaling (with exponent 0.58) of the carrying capacity with the bottleneck size. **(F)** The extinction probability of the overall *E. coli* population is plotted versus the bottleneck size (blue line), see Methods. Markers are shown for the estimated bottleneck sizes associated with SPF mice infected with two different strains, see **Figures S19B and S19C**.

Interestingly, the long-term stability differed markedly among the ESBL strains (**Figure 3**). EcST73 was particularly stable, showing no loss of bacterial colonization in any mouse (**Figures 3 and S17A**). In contrast, the other strains were comparatively less stable, with extinction observed in 20-60% of mice by the end of the experiment (**Figure 3**). Inflammation favors intestinal colonization by *Enterobacteriaceae,* especially pathobionts such as ESBL *E. coli*^38,39^. However, the lipocalin-2 (LCN2) concentration, a marker of inflammation in the feces^40^, remained low in mice colonized by each of the four strains (**Figure S17B**), suggesting that other ecological factors influenced population size and probability of extinction.

To better understand population dynamics under these conditions, we compared bottleneck sizes in LCM and SPF mice. For this we followed populations of genetically tagged strains diluted in untagged wild-type strains (**Figure S18**). From these experimental data we estimated a bottleneck (B) of 3.7×10^5^ bacteria in LCM mice colonized by EcST131 I that decreased to only 8.9 bacteria in SPF mice (**Figure S19A-B**). The intestinal load of EcST73 in SPF mice was slightly higher than that of the other strains (**Figure S17A**) and, accordingly, its estimated bottleneck size was also larger (B=39.6) (**Figure S19C**), showing that the bottleneck size scaled with the carrying capacity (**Figure 3E**). Mathematical modeling showed that the higher stability of EcST73 compared to other strains in SPF mice could be explained by its slightly higher bottleneck size **(Figure 3F)**.

Next, to investigate the mechanisms underlying ESBL *E. coli* adaptation in the gut of SPF mice, we performed whole-genome sequencing of randomly selected clones isolated from each mouse monthly. All strains accumulated mutations at a steady pace over the year (**Figure 4A**), with overall mutation rates consistent with previous reports of *E. coli* evolution in the human gut^37,41,42^ (**Figure S20A**). Across all sequenced evolved clones, we identified a total of 802 mutations, including single-nucleotide polymorphisms (SNPs), small insertions and deletions (Indels), transpositions, large rearrangements, namely inversions and large deletions (>1kb) (**Supplementary data file S2**). In three strains (EcST131 I, EcST131 J and EcST73), SNPs represented 50-70% of all detected mutations, consistent with previous results from studies of *E. coli in vivo* evolution^43^. However, strain EcST69 displayed a distinct mutational spectrum dominated by transposition events, with SNPs accounting for only ∼30% of all mutations (**Figure 4B**). This was consistent with the higher number of predicted IS*5* elements in the EcST69 genome (**Figure S20B**), which was also the prevalent IS involved in transposition events (**Figure S20C**). The mutational profiles of the strains, therefore, likely reflect strain-specific genomic features.

**Figure 4.**
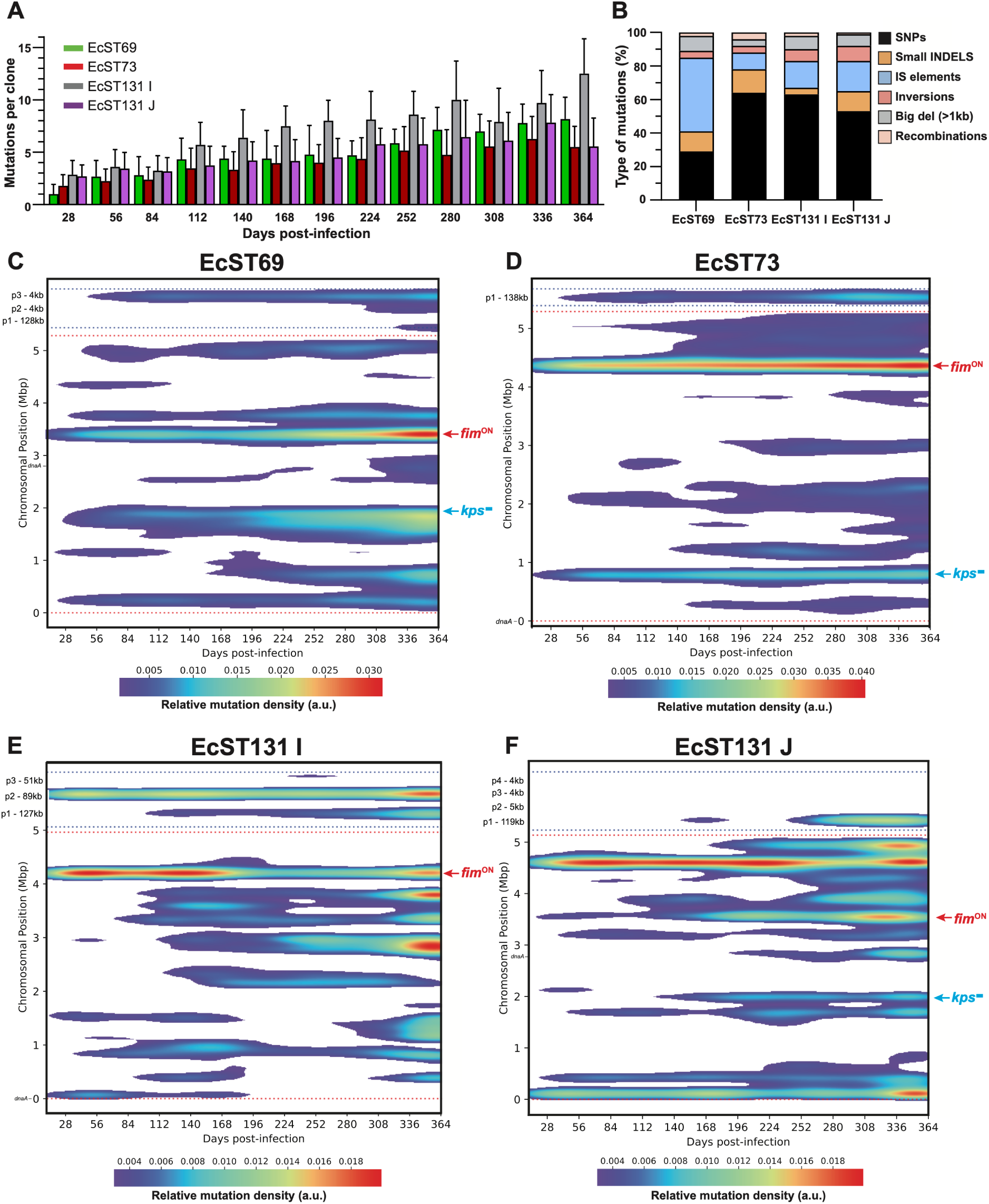
The complex gut microbiota selects for mutations involved in adhesion and biofilm formation. **(A)** ESBL *E. coli* strains accumulate mutations during gut colonization. Every 28 days, two clones of each mouse were isolated and subjected to whole-genome sequencing. The bars represent the mean and the standard deviation of the number of mutations for each clone. **(B)** The evolution of ESBL *E. coli* in the gut involves multiple types of mutation. For each strain, mutations were categorized in SNPs, small INDELs, IS elements transpositions, inversions, big deletions (over 1kb) and recombination, i.e., gene conversion. **(C-F)** Long-term gut colonization of ESBL *E. coli* is marked by convergent evolution towards *fim^ON^* and loss of capsule. The two-dimensional density maps show the distribution of mutation occurrences over time post-infection and genomic position for each strain. Mutations from all clones were pooled and genomic coordinates are shown for the chromosome and plasmids, separated by spacer regions (horizontal dashed lines). Relative mutation density (arbitrary units) reflects the local concentration of mutation occurrences after Gaussian smoothing, with higher intensity indicating regions of increased mutation clustering. *fim^ON^* and capsule mutations (*kps*^-^) are highlighted in red and blue, respectively. Details for all mutations at 364 dpi are in **Figure S21**. The details of all mutations for all time-points are in the **Supplementary data file S2**.

Analysis of non-synonymous to synonymous mutation ratios (dN/dS) revealed a spectrum ranging from neutral evolution (1) to adaptive evolution (>1) depending on the strain (**Figure S20A**). However, the majority of the mutations were observed only once, i.e., restricted to a single time point (**Figure S20A and Supplementary data file S2**), and estimations based on the rate of mutation accumulation suggested a small effective population size and a long generation time (**Figure S20A**). This indicates that non-adaptive *E. coli* evolution dominates in SPF mice, recapitulating the evolutionary dynamics of *E. coli* observed in healthy human volunteers^37^.

We nevertheless detected convergent evolution. Notably, independent lineages from all four strains acquired mutations in the *fim* operon, which encodes Type 1 fimbriae (T1F). The main target of selection was the intergenic region containing *fimS* and the *fim* promoter (**Figures 4C-F, S21, and Supplementary data file S2**). The orientation of *fimS* controls *fim* expression that can be turned ON or OFF by a single recombination event (phase variation)^44^ (**Figure 5A**). Importantly, all strains produced clones with *fimS* in the *fim^ON^* orientation, which means increased switching to the ON state, indicating selection for fimbriae expression. In addition, EcST69 acquired mutations in *fimE*, a site-specific recombinase that regulates the ON to OFF orientation of *fimS*, resulting in a *fim* “locked-ON” genotype (**Figure S21A and Supplementary data S2**). This provided a model for studying the consequences of fimbriae expression without phase variation.

**Figure 5.**
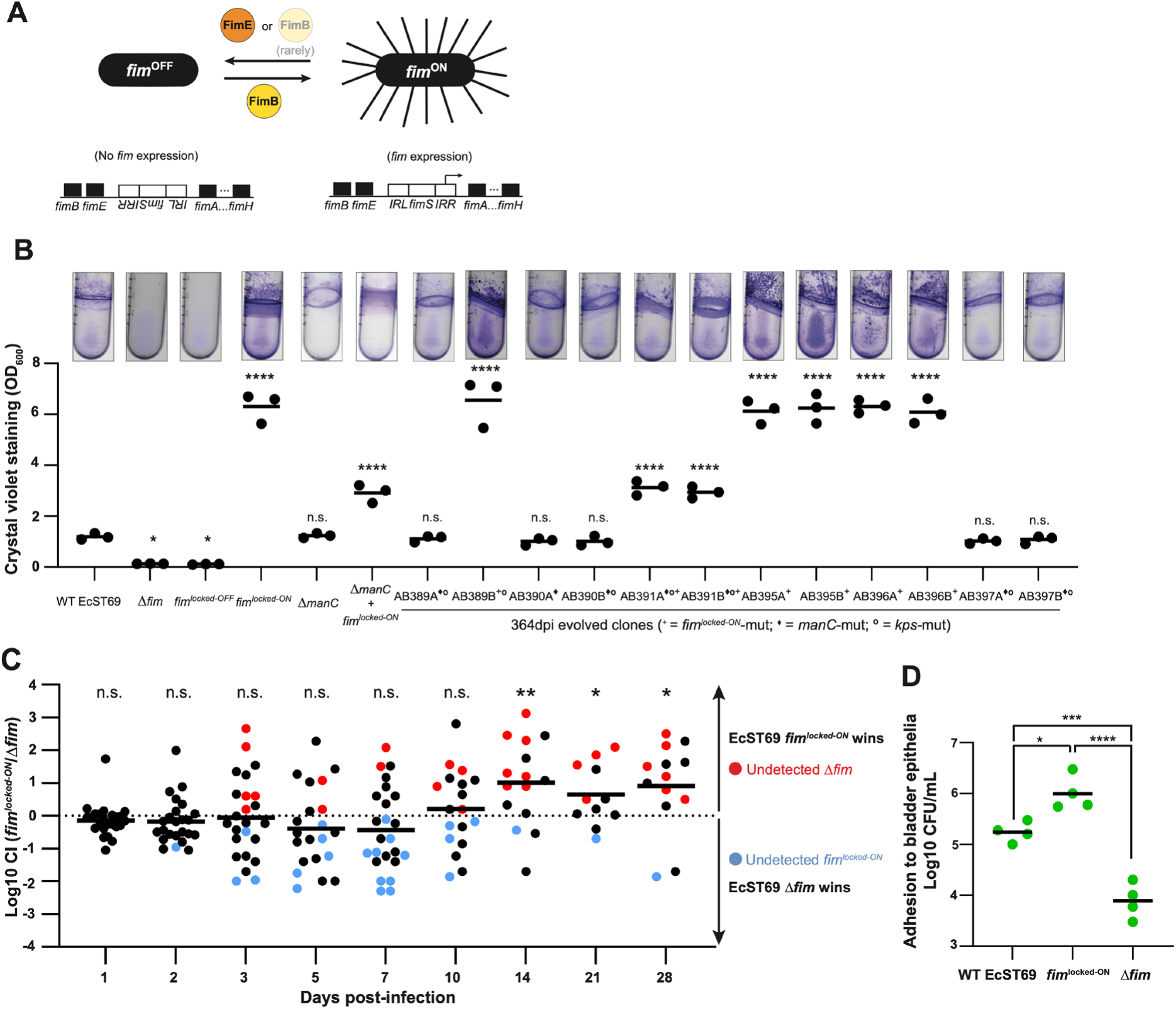
Convergent evolution towards *fim* locked-ON increases fitness in the gut and enhances pathoadaptive traits. **(A)** Schematic representation of type-1 fimbriae production controlled by phase-variation. *fim* expression depends on the orientation of the promoter region *fimS* that surrounded by the inverted repeats left (IRL) and right (IRR) and controlled by the FimE and FimB invertases. While FimE is uniquely responsible for the ON to OFF switch, FimB is able of both ON to OFF but also OFF to ON switching, although there is a preference towards the latest. **(B)** Mutations in *fimE* coupled with *fim^ON^* result in increased biofilm formation in EcST69. Top: Representative images of LB-grown cultures in 14 mL tubes for 8h after crystal violet staining; Bottom: quantification of biofilm production after crystal violet staining of cultures grown in 96-well plates for 8h in LB. The mean of three independent biological replicates is indicated by the horizontal line, and statistics were performed using one-way ANOVA with Dunn’s multiple comparisons test. Evolved clones that harbor a *fim*^locked-ON^ and/or *manC* and/or *kps* mutations are marked with the +, ◊ and ° signs respectively**. (C)** EcST69 Locked-ON *fim* outcompetes the *fim* deletion mutant in the gut. Competition assays were performed in SPF mice between *fim^ON^* and Δ*fim* for 28 days (*n*=24 mice from two independent experiments). Colored dots indicate cases in which one of the strains was not detected. Mice that lost the colonization for both strains have been excluded. Statistical tests were performed using one-sample t-test against the null hypothesis. n.s. not significant. \**p* <0.05, \*\**p* <0.01. **(D)** The Locked-ON *fim* strain shows increased adherence to human bladder tissue models. Lines represent the mean, and the statistics were performed using one-way ANOVA with Tukey’s multiple comparisons test. \**p* <0.05, \*\*\**p* <0.001, \*\*\*\**p* <0.0001.

All EcST69 clones harboring these mutations exhibited an increased adhesion phenotype quantified *in vitro* by crystal violet staining of biofilms on an abiotic surface (**Figure 5B**). The remarkable convergence of mutations affecting the *fim* system in all four strains distinguishes this as a dominant evolutionary outcome in SPF mice that confirmed the strong selection for adhesion during intestinal colonization with the microbiota. These results are consistent with our RB-TnSeq screen analyses in LCM mice, which identified strong selection for adhesion and biofilm-associated functions in the presence of microbiota (**Figures 2 and S8**).

Consistently, the reconstructed EcST69 *fim* locked-ON mutant showed a significant competitive advantage over the Δ*fim* mutant during prolonged colonization of SPF mice, demonstrating that fimbriae expression is adaptive in the gut and promotes long-term persistence (**Figure 5C**). Importantly, T1F is a crucial virulence factor that mediates adhesion to host epithelial cells, biofilm formation, and invasion of uroepithelium^45–47^. In agreement with this, the *fim* locked-ON mutant exhibited significantly enhanced adhesion in an *in vitro* human tissue model of bladder infection compared to the wild-type strain and Δ*fim* mutant (**Figure 5D**).

A second signal of convergent evolution was observed in capsule biosynthesis genes (*kps*). We found clones of EcST69, EcST73 and EcST131 J harboring loss-of-function mutations in their respective *kps* gene clusters (**Figures 4C, D, F and S21, and Supplementary data file S2**), and confirmed phenotypically by colony morphology on salt-free LB agar^48^ and/or by assessing the resistance to capsule-specific bacteriophages able to infect the ancestor strains (**Figure S22A-C**). The fact that mutations in capsule-encoding (*kps*) genes increased the fitness of EcST69 and EcST131 J in treated and in LCM mice in our RB-TnSeq screen confirmed that the loss of capsule production was adaptive during gut colonization although independently of the presence of the microbiota (**Figure S23**). This was further confirmed experimentally by a competition experiment in SPF mice, in which the reconstructed EcST69 Δ*kps* mutant showed a significant competitive advantage over the wild-type strain (**Figure S22D**).

Altogether, these findings reveal that despite genetic drift during prolonged gut colonization, microbiota-mediated competition imposes reproducible selective pressures favoring enhanced adhesion, thereby driving convergent pathoadaptive evolution in ESBL *E. coli*.

## Discussion

Intestinal carriage of ESBL *E. coli* represents a major risk factor for extra-intestinal infections (UTI, sepsis) and transmission to vulnerable hosts^2,4,6,7^. Notably, pandemic lineages of ESBL *E. coli* can persist asymptomatically in the gut for a long period without antibiotic selection, that is, in the presence of the host microbiota^19,42,49^. Therefore, fighting against these pathogens in the context of the healthy gut requires understanding how the microbiota shapes their fitness landscape and evolution. Here, we combined forward genetic screening with experimental evolution to identify the genetic determinants of successful ESBL *E. coli* intestinal colonization in the presence of a competitive microbiota. Four important findings emerge from this integrative study.

### First, we provide direct evidence that a competitive intestinal microbiota selects for diverse functions beyond nutritional competence in *E. coli* (Figures 2, S7, and S8)

The resident microbiota in LCM mice selected for specific metabolic traits in *E. coli* and, remarkably, against functions favored in its absence (e.g., xylose and maltodextrin utilization). This trade-off may contribute to maintaining metabolic versatility in *E. coli* when the pressure from the microbiota relaxes during diet fluctuations^50^, transient inflammation^38^ or antibiotic treatments^51^. In addition, the microbiota transforms the gut into a more challenging environment, in which *E. coli* fitness depends on a much broader set of functions than in the absence of microbiota. Notably, we observed a significant enrichment for pathoadaptive traits including resistance to biotic stress, cell adhesion and biofilm formation. This suggested that the competition against the gut microbiota could prime ExPEC infectivity and virulence. Indeed, the probability of causing UTI indeed increases with *E. coli’s* ability to form biofilms, notably on abiotic surfaces like catheters, to adhere tightly to the urothelium overcoming flow and stretch insults, and to survive the host immune response^6,52^.

### Second, genes that improved fitness in the presence of the microbiota were significantly overrepresented in the accessory genome (Figure S13) and some of these accessory genes were particularly prevalent in epidemic clones and known virulence factors (Figures S12 and S14)

In addition, we found two ST131-specific gene clusters that are a prophage in ST131 O25b clones and a Genomic Island (GI) present in both O25b and O16 clones. The GI harboured at least two antiphage defence systems (Lamassu and Retron) (**Figure S14**). It is unclear how these clusters could increase the fitness of *E. coli* ST131 during intestinal colonization in the presence of the microbiota. Alternatively, disrupting genes in prophages or GI encoding abortive infection systems could be slightly toxic to the cell, and this toxicity could only be detected under particularly stressful conditions, such as in the presence of the microbiota. More interestingly, we found that the siderophore Yersiniabactin encoding genes *irp1* and *irp2* in the equivalent of the High Pathogenicity Island (HPI) in our strains increased fitness when iron was limited by the microbiota^53^. Moreover, two genotoxic virulence factors in EcST73, the Cytolethal Distending Toxin (CDT) and the colibactin were positively selected. CDT could promote interaction with epithelial cells by disrupting their cell cycle^54^ and colibactin could contribute to eliminating competitors in the gut^55,56^. Hence, competition with the microbiota, but not the growth in the gut *per se*, selected for oncogenic traits in *E. coli*. Moreover, several genes that increase fitness in the presence of microbiota were not detected in *E. coli* genomes from B2 phylogroup despite being broadly distributed across other phylogroups (**Figure S14**). This uneven distribution of fitness determinants suggests that distinct phylogroup lineages may rely on different sets of genes for colonizing the competitive intestinal environment.

### Third, ecological constraints in a complex microbiota destabilize *E. coli* populations

As expected in a constrained metabolic landscape, the host carrying capacity (K) for *E. coli* was limited by microbiota complexity (**Figures 3, S2, and S18**). Moreover, *E. coli* experience population bottlenecks (B) of a size that depends on the complexity of the microbiota (**Figures 3, S18, and S19**). The conventional microbiota in SPF mice maximizes constraints, thereby drastically limiting the population size of ESBL *E. coli* (**Figure 3**). By comparing treated (high K), LCM (intermediate K) and SFP mice (low K), we found that the size of B scales with K, meaning smaller bottlenecks at lower K (**Figures 3E, S18 and S19**). Accordingly, the effect of disrupting biofilm formation on the relative fitness of EcST73 was amplified in SPF compared to LCM mice, and effectively lost in antibiotic-treated LCM mice (**Figure S24**). This suggests that traits that increase the probability of surviving repeated narrow bottlenecks are particularly relevant in the presence of a complex microbiota.

Modelling also explains the remarkable stability of EcST73 in SPF mice that correlates with slightly increased K and B compared to the other strains in this context (**Figures 3E, 3F, and S19**). The model indeed shows that such a slight increase in a low K is sufficient to make population extinction extremely unlikely (**Figure 3F**). Correspondingly, very low K and narrower bottlenecks could explain the occasional extinction of the other strains in SPF mice (**Figure 3F**). More work is required to identify the underlying mechanism of EcST73 stability in SPF mice. However, these observations in our conventional mouse line, which did not harbour any other *Enterobacteriaceae*, recapitulated the relative instability of ESBL *E. coli* in the intestinal tract of healthy humans^57,58^, where the effective population size of *E. coli* is small^37^. Narrow bottlenecks in the presence of a complex microbiota generate stochastic extinction, thereby ruling out the necessity of competing *E. coli* strains to exclude ESBL *E. coli* from healthy carriers.

### Fourth, phase variation offsets the dominance of neutral evolution in the presence of a complex microbiota

The relative stability of the ESBL *E. coli* strains in SPF mice enabled us to follow their evolution in the presence of a complex microbiota over the course of a year. *E. coli* colonization of SPF mice recapitulated the dominance of genetic drift observed when *E. coli* evolves in the gut of healthy humans that results from a small effective population size^37^ (**Figure S20**). Nevertheless, adaptation by increasing adhesion occurred and was convergent across the four strains. This was possible because the expression of the Type 1 fimbriae (T1F, *fim* operon) can be turned ON at high frequency via a site-specific recombination event (phase variation^44^) in a substantial fraction of fecal clones. In EcST69, additional mutations locked the expression of the *fim* operon via inactivation of *fimE*. The expression of T1F increased *in vitro* biofilm formation in EcST69, which was not observed for the other strains. This may be due to mannose residues in the O17 antigen^59^ of EcST69 that T1F can bind to, thereby fostering auto-aggregation. Apart from gut colonization, T1F locked-ON expression increased attachment to a human urothelial tissue model, which agrees with the critical role of FimH adhesin during bladder colonization^52,60^ (**Figure 5D**). Interestingly, some EcST69 fecal clones harboured additional mutations in *manC,* resulting in a short O-antigen (**Figure S22E**). The reduced ability of these mutants to form biofilms compared to locked ON T1F strains (**Figure 5B**) suggested progressive fine-tuning of stickiness during within-host evolution.

**In conclusion**, the genetic basis of *E. coli* fitness in the gut is strongly contingent on the microbial context. Interactions with the resident microbiota define complex colonization strategies that involve more than optimizing the use of limited resources to grow. Instead, ecological competition during asymptomatic gut colonization selects for pathoadaptive traits, including adhesion, biofilm formation, and diverse accessory virulence genes. Thereby, this work demonstrates that the gut is not just a reservoir but is also a niche that primes ESBL *E. coli* for extra-intestinal virulence.

## Acknowledgments

We thank to the member of the Diard lab for the discussion; the staff at the core and animal facilities at the University of Basel and ETH Zürich for their support; Bidong D Nguyen, Benjamin B. J. Daniel, Yves Steiger, Julia A. Vorholt, and Wolf-Dietrich Hardt for sharing the RB-TnSeq plasmid, primer details, and WISH-tag plasmids; Lena Siewert and Anne-Katrin Pröbstel for their help with germ-free mice; Alexander Harms for sharing *E. coli* strain MG1655; Luca Wolf, Alexander Wenzel, Rahel Kocher, Louna Tscheulin, Pauline Schamberger, Jerrik Batram, Victor Cappoen, and Valentin Druelle for technical assistance.

## Author contributions

Conceptualization: P.K.J., L.R., and M.D. Methodology: P.K.J., L.R., M.M., N.W., A.S., S.S., A.F.B., and M.D. Investigation: P.K.J., L.R., M.M., N.W., M.M., C.F., C.F., R.M., L.D., A.B., S.F., E.L., M.J., M.K., and A.R. Visualization: P.K.J., L.R., P.M., M.M., C.F., A.S., A.F.B. and M.D. Funding acquisition: M.D. Writing - original draft: P.K.J., L.R., and M.D. Writing - review and editing: P.K.J., L.R., M.M., N.W., C.F., C.F., P.M., C.F., A.S., R.M., L.D., A.B., S.F., E.L., M.J., M.K., A.R., A.E., U.J., C.D., S.S., A.F.B., and M.D.

## Data and materials availability

Codes for RB-TnSeq data analysis are available https://github.com/MicrobiologyETHZ/mbarq and https://github.com/MicrobiologyETHZ/pjangir_rbtnseq. Barcode mapping and barcode counts files are available from GitHub https://github.com/MicrobiologyETHZ/pjangir_rbtnseq. Codes for the analysis of *E. coli* genomes are available from GitHub https://github.com/mmolari/microbiota-genes.

## Supplementary materials

### MATERIALS AND METHODS

The list of chemicals, commercial kits, and other materials can be found in Table S1.

#### Animals

All experiments performed with mice followed the 3Rs guidelines to reduce animal use and suffering, and were approved by the legal authorities (licence number: 2987_36493, Kantonales Veterinäramt Basel-Stadt, Switzerland). All breading and housing conditions consisted of individually ventilated cages with a 12-hour light/dark cycle, constant ambient temperature (22 °C + /− 1°C) and humidity (60 % − 40 %) and with an *ad libitum* standard chow diet either in the ETH Zurich Phenomics Center (EPIC) or at the Werk Rosental Animal Facility of the University of Basel (WRO). Eight- to twelve-week-old male and female C57BL/6J mice were randomly assigned to each experimental group in equal numbers, whenever possible. Three different types of mice were used (i) conventional Specific Pathogen Free (SPF), containing an intact and complex intestinal microbiota with a high colonization resistance, (ii) LCM (Low Complexity Microbiota), characterized by a decreased colonization resistance that facilitates *E. coli* colonization^1,2^, and (iii) germ-free (GF) (TCR MOG heterozygous line on the C57BI/6J background) mice. For intestinal microbiota depletion by antibiotic treatment, a single dose of ampicillin (20 mg per mouse) was administered by oral gavage 24 h prior to infection, followed by ampicillin supplementation in the drinking water (2g/L) 24 h after the first administration. Depending on the experiment, animals were either co-housed or single-housed to prevent transmission events. Prior to infection, feces were collected and stored for further analysis, and to confirm the absence of any contamination. Of note, all mouse models used in this study are naturally *E. coli*-free. This was routinely confirmed by plating the homogenized feces on pure MacConkey (Oxoid) agar plates to select for *E. coli* (see conditions below).

#### Strains, media and growth conditions

The bacterial strains, plasmids and medium supplements used in this study are provided in the Table S1. The four ESBL *E. coli* isolates used throughout this project were isolated from healthy human volunteers and were subjected to whole-genome sequencing. Unless mentioned otherwise, all liquid bacterial cultures were grown aerobically in LB (10 g/L tryptone, 5 g/L yeast extract, 10 g/L NaCl) with agitation (200 RPM) at 37°C. An overnight culture corresponded to 16-20 h of growth. For the preparation of electrocompetent ESBL *E. coli*, bacteria were grown to exponential phase (OD_600_ 0.5-1) at 30°C in salt-free LB (10 g/L Tryptone, 5 g/L yeast extract) and washed twice with sterile cold water. Growth on solid LB agar (LB+1.5% agar) or on MacConkey agar plates was also assessed at 37°C. When required, antibiotics were added to the media: ampicillin (Amp) 100 μg/mL, chloramphenicol (Cm) 15 μg/mL, and kanamycin (Km) 50 μg/mL. Gentamicin (Gm) was used at 10 μg/mL in low-salt LB (10 g/L tryptone, 5 g/L yeast extract, 5 g/L NaCl). To grow *E. coli* JKe201^3^, 1 mM of 2,6-Diaminopimelic acid was added to the medium.

##### Construction of randomly barcoded transposon mutant libraries (RB-TnSeq libraries)

RB-TnSeq mutant libraries were constructed as previously described^4^. Briefly, the DNA-barcoded EZ-Tn*5* transposome was constructed by PCR from the linearized (NotI-HF digested) plasmid template pZ922 (a generous gift from Prof Wolf-Dietrich Hardt’s lab, ETH Zürich). PCR amplification of the kanamycin resistance cassette was performed using primers that introduced a randomized 17-nucleotide barcode sequence and the Tn*5* mosaic end inverted repeats. Primer tn5+ME+17N-F3 incorporated the 17N barcode adjacent to the RB-TnSeq priming region, whereas primer tn5+ME-R2 completed the mosaic end structure. PCR reactions were carried out in 100 μl volumes containing ∼40 ng of digested plasmid DNA template, 0.1 μmol of each primer, and 0.5 μl Phusion DNA polymerase. Cycling parameters included an initial denaturation at 94°C for 1 min, followed by 30 amplification cycles (94°C for 30 sec, 55°C for 30 sec, and 72°C for 1 min), and a final extension at 72°C for 5 min. The amplicon was purified from a 1% agarose gel using the ReliaPrep-Kit, and eluted in TE buffer. For transposome formation, 500 ng of purified transposon DNA was mixed with 2 μl of EZ-TN5 Transposase and 1 μl of 100% glycerol. After brief mixing, the mixture was incubated for 1h at 37°C. To remove excess salt impurities, drop dialysis on a Type-VS Millipore membrane (0.025 μm pore size) was performed for 30 min at room temperature. Multiple reactions were prepared to generate a high-density and a diverse mutant library, which was further validated by sequencing (see methods below). Approximately 200-300 ng of the DNA-barcoded EZ:Tn*5* transposome was electroporated into 60 μl of freshly prepared electrocompetent *E. coli* cells. Cells were immediately recovered in SOC medium, incubated for 1h at 37°C with shaking, and plated in batches on LB agar (50 μg/mL kanamycin). The transposon mutants were recovered the following day, and aliquots of a pool of approximately 5000 colonies were stored in LB+glycerol at −80°C.

##### Sequencing library preparation and mapping of the transposon insertion sites

To sequence the constructed RB-TnSeq libraries, genomic DNA was isolated separately from each library pool (containing ∼5000 mutants) using the Quick-DNA Miniprep Plus Kit (Zymo). The purified high-quality genomic DNA (∼1 µg) transferred to covaris tubes (8 AFA-TUBE TPX) and subjected to shearing using Covaris sonication machine with the following parameters: base pair mode (bp): 175 – 350; sample volume (µL): 20 – 50; temperature (°C): 10; peak incident power (W): 220; duty factor(%) = 25; cycles per bust (cpb): 50. The empty wells in the tubes were filled with the same volume of TE buffer. The desired fragment size was 300-500bp. The fragment size and quality of the sheared DNA were confirmed using the Agilent TapeStation system. The sheared DNA was used to prepare a sequencing library using the NEBNext Ultra II DNA Library Prep Kit (Illumina) with a minor modification as follows. For the PCR enrichment of adaptor-ligated DNA, we used a custom made i5* primer which binds to the transposon, ensuring the enrichment of transposon containing DNA. PCR cycling parameters included an initial denaturation at 98°C for 30 sec, followed by 25 amplification cycles (98°C for 10 sec and 65°C for 75 sec), and a final extension at 65°C for 5 min. The amplicon was purified using AMPure beads as described in NEBNext Ultra II DNA Library Prep Kit (Illumina). Illumina paired-end sequencing with 150 bp read length was performed.

To map the insertion site of each barcoded transposon, reads from the sequenced library pools were processed and analysed using a previously established mBARq tool with default parameters^5^. Briefly, only reads with a perfect match of at least 15 base pairs to the Tn*5* transposon end sequence were selected for subsequent analysis. The filtered transposon reads were aligned onto the genome sequence of the corresponding wild-type (WT) *E. coli*. Barcodes were annotated to a specific gene if the insertion was found within the protein-coding sequence, and unannotated barcodes were excluded for the downstream analysis. Intergenic insertions were also identified and included in the data analysis.

##### *In vivo* screening of RB-TnSeq libraries

RB-TnSeq libraries were screened in the presence of microbiota (LCM mice) and in the absence of microbiota (antibiotic-treated LCM and germ-free mice). Frozen library aliquots, each containing approximately 5000 distinct mutants, were thawed on ice and incubated in LB broth at room temperature for approximately 30 min. Individual aliquots were pooled to generate a combined library targeting ∼30000 different barcoded mutants, with minor variation depending on the library size of the strain. The pooled library was washed twice with sterile phosphate-buffered saline (PBS) and resuspended in 100 µL of PBS buffer. This final inoculum contained ∼10^9–10^ CFUs (colony forming units) in total and was administered via oral gavage to mice (100 µL dose per mouse), ensuring multiple cells per mutant. To define the composition of the input (before infection) library pool, two aliquots of the pooled library were used as inoculum replicates. For each library, 8 mice were used as biological replicates post infection, with each mouse housed in a separate cage to prevent any transmission. Fresh fecal pellets were collected from each mouse on day 1, 2, 3, 4, and 8 post-infections. Mice were sacrificed on day 8 post-infection using CO_2_ after collecting the last time-point sample.

Feces pellets were homogenized (25Hz for 3 min with 3mm glass beads) in 1 mL of PBS using the Tissue Lyser device (Qiagen), and serial dilutions were plated on LB kanamycin plates to determine total bacterial counts. To enrich the RB-TnSeq mutants in feces samples, 700 µL homogenate was incubated for 4h at 37°C with shaking in 15 mL of LB containing kanamycin (50 µg/mL). The same enrichment procedure was applied to the two input replicates. After enrichment, cells were harvested by centrifugation at 4500 RPM at room temperature, and resuspended in 150 µL of TE buffer and stored at −20°C until further processing for barcode sequencing and counting.

##### Barcode counting and fitness calculation

Genomic DNA was extracted from enriched RB-TnSeq mutants using the QIAamp Fast DNA Stool mini kit (Qiagen) according to the manufacturer’s instructions with the following modifications: 900 µL supernatant from step 4, adding 650 µL Buffer AL at step 7 and 750 µL of ethanol to the lysate at step 9. Elution was performed in 40 µL of ATE elution buffer. The barcode region was PCR-amplified using primers that enable sample multiplexing (see Table S1). For each mouse sample, a unique index was incorporated using the indexing primers. The following cycling program was used: 98°C for 30s, 28 cycles of 98°C for 10s, 60°C for 30s, 72°C for 20s, followed by a final elongation at 72°C for 5 min. PCR products were purified using the ReliaPrep^Tm^ DNA Clean-Up and Concentration kit (Promega) and stored at −20°C. Up to 14 samples were pooled equimolar (500 ng each) and gel-purified from a 1.7% agarose gel using the ReliaPrep^Tm^ DNA Clean-Up and Concentration kit. The purified pools were sent for paired-end NovaSeq150 Illumina sequencing at Novogene (UK). Samples with insufficient DNA yield and sequencing quality were excluded from downstream analyses (e.g., AB963 D1 and AC069 D8). In addition, no fecal sample was available for AB968 D8.

The RB-TnSeq sequencing files were demultiplexed, and the abundance of each barcode in each sample was calculated using the previously established mBARq tool^5,6^ (mbarq count command with default parameters). In the resulting barcode counts file, barcodes with fewer than 100 reads across all samples (two input replicates and eight mouse samples across time points) were removed from further analysis. This filtering step helps to reduce noise from barcodes with very low read counts, which may introduce variability in downstream analysis. To enable comparisons across experimental conditions (LCM, antibiotic-treated, and GF), a filtering step was implemented to retain only barcodes detected in at least one sample from each condition. This approach ensures that downstream analyses compare identical mutant populations across different microbiota environments. Each strain was analysed independently. Additionally, to ensure robust fitness measurements, only samples showing reproducibility between replicates (Pearson correlation >0.6) at the barcode counts level were included in the fitness analysis.

Next, we identified differentially abundant mutants between the control (the inoculum) and the treatment conditions using the mbarq tool (mbarq analyse command with default parameters)^5^. For this fitness calculation and comparative analyses between conditions or strains, only early infection time points were included that showed stable retention of genomic features and maximized infection duration (**Figure S3**), while avoiding effects of population bottleneck and/or clonal expansion. Accordingly, day 1 post-infection fitness data were used for EcST131 I and EcST73, and day 2 post-infection fitness data were used for EcST131 J and EcST69. Genes showing at least 2-fold increase or decrease in fitness (i.e., log2 LFC ≥ 1 or ≤ −1) with an FDR <0.05 were considered significant hits. The RB-TnSeq fitness data are provided in the **Supplementary data file S1**.

##### Principal component analysis

To enable comparisons across experimental conditions (LCM, antibiotic-treated, and GF), a filtering step was implemented to retain only barcodes detected in at least one sample from each condition. This approach ensures that downstream analyses compare identical mutant populations across different microbiota environments. Each strain was analysed independently. Barcode counts were normalized using a log₂ counts-per-million (log2 CPM) transformation. For each sample, raw counts were first converted to counts per million by dividing by the total library size and multiplying by 10⁶. The resulting values were then log₂-transformed with a pseudocount of 0.5 to handle zero counts. Principal Component Analysis (PCA) was performed on the normalized count matrices to assess global patterns of population structure across samples. The analysis used the scikit-learn implementation (v.1.5.1) with default parameters^7^, extracting the first two principal components. All analyses were implemented in Python 3.9 using the following libraries^8^: pandas (v.2.3.3) for data manipulation^9^, NumPy for numerical operations (v.1.26.4)^10^, scikit-learn (v.1.5.1) for PCA, and Plotly (v.5.24) for visualization^11^.

##### String network and Gene ontology (GO) enrichment analysis

Genes showing a significant fitness change were analyzed using the STRING protein-protein interaction database to identify functional gene clusters^12,13^. For string network figures, genes were grouped into manually curated functional categories based on GO annotations and established gene functions (see the Source data file). GO enrichment analysis was performed to determine biological processes overrepresented among genes increasing fitness specifically in the presence of microbiota or in the absence of the microbiota. GO annotations were obtained from the GO database. GO categories were selected based on functional clusters identified in the STRING protein-protein interaction network. Enrichment was assessed using Fisher’s exact test, and p values were adjusted for multiple testing using Benjamini–Hochberg false discovery rate (FDR) method in R^14^.

##### Enrichment of core and accessory gene

To investigate whether genes increasing fitness specifically in the presence of microbiota were enriched within the core or accessory genome, all *E. coli* genes were first classified as either core or accessory. For this, we analyzed 51 publicly available genomes, including representatives from prevalent sequence types (such as ST131, ST69, ST73 as well as common laboratory strains (e.g., *E. coli* strain MG1655 and BW25113), to assess whether genes of interest were also present in lab-adapted backgrounds (**Supplementary data file S3**). Genes from these isolates were clustered using PanX^15^, and gene clusters present in at least 95% of the 51 genomes and not duplicated in any genome were classified as core, and the rest as accessory. Enrichment among core or accessory genes was assessed using Fisher’s exact test.

##### Core-genome phylogeny tree

To construct the core-genome phylogeny we used the collection of diverse 51 *E. coli* genomes used for the enrichment of core and accessory gene analysis. We then queried RefSeq^16^ for all complete assemblies of taxon “escherichia coli”, and selected a random subset of 300 elements with “Homo Sapiens” as a host. The final collection comprises 352 genomes. Using PanX^15^ we performed gene clustering on the full collection to identify core genes, and concatenated their alignments to create a core-genome alignment. From this alignment we generated a core-genome phylogenetic tree using FastTree^17^ and refined branch lengths by maximum-likelihood optimization using TreeTime^18^.

##### Genome collection and comparative pangenome analysis

###### The Horesh *E. coli* collection

To investigate how our results generalize to the broader *E. coli* species, we analysed the genome collection introduced in a recent study (referred as the Horesh collection)^19^. The collection consists of more than 10′000 curated *E. coli* and Shigella assemblies, complete with relevant metadata. The authors performed gene-clustering on the 50 largest lineages (∼7000 genomes) and provide a pangenome reference and a binary gene presence/absence matrix for these genomes. We restrict our analysis to these isolates. Finally, a core-genome tree is provided that includes 500 representative genomes. Amongst other metadata, genomes in the collection have been assigned a putative sequence type (ST) and phylogroup. The ST distribution is skewed, with the eight largest STs alone accounting for ∼ 73% of the dataset (ST11, *n* = 3050; ST131, *n* = 761; ST73, *n* = 431; ST95, *n* = 286; ST21, *n* =209; ST17, *n* = 184; ST69, *n* = 163; ST442, *n* = 124). But in total we find 55 STs with ≥ 10 isolates.

###### Identification of the fitness-increasing (microbiota-specific) genes in the Horesh collection

In our RB-TnSeq experiments we identified a set of 506 genes that increase *E. coli* fitness in the presence of microbiota. We used nucleotide alignment to identify these genes in the Horesh collection. We mapped the consensus sequences of these genes, as extracted from our PanX gene clustering, to the pangenome reference in the collection using nucleotide BLAST, retaining hits with percent identity ≥ 85%, query coverage ≥ 85% and subject coverage ≥ 85%. We found a 1-1 mapping for the majority of our candidates (*n* = 436). For *n* = 49 we found multiple hits, which were subsequently merged into a single cluster. For *n* = 21 genes we could find no correspondence within the desired similarity in the Horesh pangenome, and they were subsequently dropped. This results in a restricted pangenome presence/absence matrix for 485 genes over 7512 genomes, that is used as input for our subsequent analysis.

###### High-load strains

Genomes have on average *n* = 308 ± 23 genes (those that increase fitness in the presence of microbiota), with systematic difference between different STs, see **Figure S12A**. We define high-load strains as the ones with *n* ≥ 330 genes. This partitions the collection into 1′878 high-load and 5′634 normal-load strains.

###### ST participation ratio

For each of the gene that increase fitness in the presence of microbiota, we calculate its ST participation ratio as: 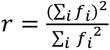, where *f_i_* is the empirical frequency of the gene in ST *i*, i.e. the fraction of genomes in ST *i* in which the gene is present. This is calculated over all STs with *n* ≥ 10 genomes. The number *r* quantifies the effective number of STs in which the gene is commonly found.

###### Pairwise phylogenetic distances

To compute phylogenetic distances between STs we rely on the core-genome tree provided with the dataset. The tree contains 500 representative genomes. We restrict the analysis to the 51 STs that have at least 10 representatives in the dataset, and at least 1 representative in the tree.ST average distances are calculated by averaging the tip-to-tip distance in the tree between all unique pairs of genomes from the two considered STs.

##### Genome-editing techniques

All the strains used in this study are listed in Table S1. All the plasmids constructed in this study are derivatives of the published plasmids pFOK^20^, pEMG^21^ and pXG10-SF^22^ and are listed in Table S1. Gene deletions were performed by *λ red* recombination^23^. ESBL *E. coli* strains that carried the helper plasmids pSIM5^24^ or pSIM5-*tet*^25^ were grown in salt-free LB (without antibiotics) at 30°C until the exponential phase was reached. The *λ red* genes were induced at 42°C for 45 min, and bacteria were washed twice with cold sterile water. The bacteria were concentrated 100 times in water, and 50 µL of the resulting competent cells were electroporated with 0.5-2 µg of PCR fragments that carried antibiotic resistance genes flanked by *frt* sequences and 40 nucleotides, that corresponded to the chromosomal targeted region. The PCR fragments were amplified with Phusion DNA polymerase, specific oligonucleotide pairs (see Table S1) and the template plasmids, pKD3 (Cm^R^), pKD4 (Km^R^) or pKD4-Gm, a pKD4-derived plasmid carrying the *aacC1* gentamicin resistance gene. After selection with the appropriate antibiotic, recombinant ESBL *E. coli* were transformed with the FRT-flippase encoding plasmids pCP20^26^ or pCP20-TcR^27^ to eliminate the antibiotic resistance cassette. Temperature-sensitive plasmids pSIM5, pSIM5-*tet*, pCP20 or pCP20-TcR were eliminated by a passage at 42°C. All the mutations were confirmed by PCR and Sanger sequencing (Microsynth AG, Balgach, Switzerland).

To generate scarless *malT* deletion in EcST73, a suicide plasmid-based allelic exchange approach was used^20^. The **∼**750 bp regions flanking *malT* were PCR-amplified using the primer pairs OMD25_348/OMD25_349 and OMD25_ 350/OMD25_ 351. The Two PCR amplicons were fused by overlap-extension PCR. The resulting fragment was digested with *Bam*HI and *Not*I and was ligated (T4 DNA ligase) into the corresponding sites of the suicide plasmid pFOK. The resulting plasmid was introduced into *E. coli* JKe201 and was mobilized by conjugation into the EcST73 strains of interest. The EcST73 recombinants were selected on LB agar plates supplemented with kanamycin, and merodiploids were resolved as described by Cianfanelli and colleagues^20^.

The *fim* locked-ON strains (mutation Δ*fimBE fimS^ON^*) were generated by a two-step *λ red* recombination protocol. For EcST69 the *frt-aphT-frt* module of pKD4 was PCR-amplified with primers OMD25_326 and OMD25_027 and recombined into an *in vivo* evolved strain carrying a spontaneous *fim* locked-ON mutation (MDBZ2905). The *frt-aphT-frt-fimS^ON^* module of the resulting strain was PCR-amplified with primers OMD25_326 and OMD25_042 and the PCR fragment was recombined into EcST69.

For the chromosomal tagging of the resulting strains, a Tn*7*-based tagging method^28^ was used to introduce the *cat* (Cm^R^) and the *aphT* (Km^R^) resistance genes into the *glmS-pstS* locus, using the pGRG36 Tn*7* delivery plasmid^29^.

The laboratory strain *E. coli* MG1655 used for the SPF mouse colonization experiment carried a restored O-antigen (*wbbL*+, functional O-16)^30^. The strain was tagged with the *aphT* (Km^R^) marker using Tn*7*-based chromosomal tagging, and carried the pXG10*-oriT-traJ* plasmid.

##### Microbiota analysis

Fecal samples from uninfected SPF and LCM mice were sampled, processed, and stored in glycerol as described above. Genomic DNA isolation, 16S rRNA gene amplification and sequencing were performed by Zymo Research Europe using the 16S Targeted Sequencing service. Briefly, DNA was isolated from feces using the ZymoBIOMICS-96 MagBead DNA Kit (Zymo Research, Irvine, CA) and the targeted library preparation was performed with the Quick-16S™ Primer Set V3-V4 (Zymo Research Europe, Freiburg, Germany). Sequencing was carried out on the Illumina MiSeq platform. Unique amplicon sequences were inferred using the Dada2 pipeline^31^ and the taxonomy assignment was performed using Uclust from Qiime (v.1.9.1)^32^ with the Zymo Research Database, an internally curated 16S database, as the reference.

##### Long-term *in vivo* evolution experiment (LTEE)

Twelve conventional SPF mice were co-housed, three animals in each cage, and monitored on a weekly basis over one year. Each cage had a different inoculum derived from an independent overnight culture. For the inoculum preparation, the corresponding *E. coli* strain was grown overnight (from the glycerol stock) in 10 mL of LB medium supplemented with ampicillin (100 μg/mL) at 37°C with shaking. For each mouse, 1 mL of the overnight grown bacterial culture was centrifuged at 10,000 *g* for 3 min at room temperature, and the bacterial pellet was washed twice with 1 mL of PBS buffer. The washed bacterial pellet was then resuspended in 100 µL of PBS buffer. Mice were infected by oral gavage with the entire volume of the inoculum (100 µL), which resulted in approximately 10^9^ CFUs. To measure the inoculum size, CFU counting was performed by serial dilutions in PBS buffer of an extra inoculum and spotting on MacConkey+ampicillin (100 μg/mL) plates. For each sampling day, feces samples were collected into a 2 mL tube, resuspended in 1mL of sterile PBS by shaking at 25Hz for 3 min with two 3mm glass beads, then mixed with glycerol (10% final concentration) and stored at −80 °C for further analysis. The *E. coli* colonization levels were assessed by plating 100 µL of undiluted resuspended feces and/or by serial dilutions in PBS and by spotting on MacConkey+ampicillin (100 μg/mL) plates, followed by CFU counting. *E. coli* colonization was checked at 1, 2, 3, 5, 7, 10, 14, and 28 days post-infection (dpi) and then every 28 days until 364 dpi. After 182 dpi, cages in which all three mice remained non-colonized for more than 28 days were sacrificed. For EcST73 experiment, three mice (cage D) were prematurely excluded due to a technical issue and the last time-point available is the 308 dpi.

##### Whole genome sequencing

For genomic DNA extraction, bacterial cells were grown overnight in LB medium supplemented with ampicillin (100 μg/mL). Cells were harvested by centrifugation, and the genomic DNA was purified using the Quick-DNA Miniprep Plus Kit according to the manufacturer’s instructions, but with an additional RNase A (500µg/mL) treatment after the addition of the Red lysis buffer (Zymo). All original ancestral strains were sequenced using both short-read Illumina and long-read Oxford Nanopore Technologies sequencing with SeqCenter’s Small Nanopore Combo service (Pittsburgh, USA) followed by assembly (Unicycler v0.4.8^33^) and annotation (Prokka v1.14.5^34^) (https://www.seqcenter.com/services/).

For the LTEE, two clones were isolated from each colonized mouse at 28-day intervals until 364 dpi. These evolved clones were sequenced in-house using long-reads Oxford Nanopore Technologies (MinION Flow Cells R10 and Rapid Barcoding Kit 24). Basecalling was performed with Guppy (version 6.5.7+ca6d6af; dna_r10.4.1_e8.2_400bps_sup; Oxford Nanopore Technologies). To determine the type, frequency, position, and linkage of mutations over time, sequencing reads from each evolved clone were mapped to the corresponding ancestral strain and analyzed using the Evo-genome-analysis pipeline (https://github.com/mmolari/evo-genome-analysis), as previously described^35^. Detailed mutation profiling data for each strain and clone are provided in the **Supplementary data file S2**. The frequency thresholds for detecting different mutations were *consensus* > 0.9, *gaps* > 0.9, *insertions* > 0.3, and *clips* > 0.15. Inversions were considered when the clip frequency was above the refence (i.e., ancestral wild-type strain).

##### Calculation of the mutation rate, dN/dS and estimation of the generation time and effective population size

The mutation rate was calculated using the formula 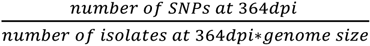. All SNPs identified at 364 dpi were included in this calculation, encompassing coding and non-coding (intergenic) regions, as well as plasmid sequences. The dN/dS ratio was calculated considering only SNPs located in coding regions at 364 dpi. One-time mutations were defined as mutations detected only once (i.e., in one or two clones from the same mouse or different mice at the same time-point) and not detected in subsequent samples. Generation time was estimated based on a previously described method^36^, using the formula 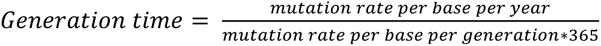, where only chromosomal mutations were considered for the analysis. The extreme estimates for the mutation rate per base per generation in the gut were 0.89×10^−10^ and 3.12×10^−10^. The effective population size was estimated as previously described^36^, using the formula 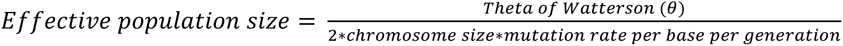. In our case, for each such pair of clones at the same time-point, we can calculate the Watterson θ estimator, which in this case is effectively equivalent to the number of single-nucleotide differences between the two considered clones sampled from the same mouse for the same time-point. In the hypothesis that the effective population size and mutation rate is roughly stable across mice and timepoints for the same strain, then we expect the number of SNPs between such pairs of clones to be geometrically distributed. In this case, the maximum-likelihood estimate for the θ for a specific strain is given by the average number of SNPs between all co-sampled clone pairs (isolated from the same mouse) belonging to the same strain.

##### 2-dimensional density plots for displaying the mutation occurrences over time during the LTEEs

To visualize mutation kinetics during long-term intestinal colonization, mutation events from clones isolated from all mice and cages of the same strain were combined into a single dataset. Each mutation was mapped to a genomic coordinate according to its chromosomal or plasmid location. For mutations annotated as genomic intervals, the midpoint of the interval was used as a representative position. Mutation density was computed as a two-dimensional histogram of mutation events as a function of time post-infection and genomic position. The resulting histogram was independently smoothed along the temporal and genomic axes using a Gaussian filter (scipy.ndimage.gaussian_filter) to generate a continuous density map. To improve visual interpretability, low-density regions were attenuated by applying a percentile-based threshold. All analyses and visualizations were performed in JupyterLab using NumPy (version 2.3.5), SciPy (version 1.16.3), and Matplotlib (version 3.10.6) packages.

##### Competition assays *in vivo*

For an *in vivo* competition experiment, independent inoculum cultures were prepared by growing bacterial strains in 10 mL of LB+ampicillin (100 μg/mL) medium for overnight at 37°C with shaking. After, the OD600 was measured and corrected if needed, the cultures were mixed at a 1:1 ratio. The mixture was then centrifuged (15 min, 4500 RPM, room temperature), and the bacterial pellet was washed twice with 10 mL of PBS. The bacterial pellet was centrifuged again with the same conditions and then resuspended in 1 mL of PBS. The ratio of the strains in the inoculum was measured by CFU counting on selective LB agar plates. Independently of the mouse model, each animal was infected by oral gavage with 100 µL of inoculum, which resulted in approximately 10^9^ CFU per dose. The competitive index was determined over time by CFU counting on selective LB agar plates and corrected by the initial inoculum ratio.

Competition experiments in LCM mice were restricted to a maximum of a week post-infection because higher population size in LCM mice increases the likelihood of *de novo* adaptive mutant emergence during prolonged colonization which can confound competitive fitness measurements. In particular, maltose mutants (e.g., *malT*) emerge in a week in LCM mice, depending on the strain. To further control for the potential impact of spontaneous maltose mutants, competition experiments involving the EcST73 Δ*pga*Δ*csg* double mutants were additionally performed in a Δ*malT* background, in which both the mutant and the wild-type control strain carried the *malT* deletion.

##### Quantification of biofilm formation by crystal violet staining

Three independent cultures were started from single colonies, previously streaked from the cryo-stocks and grown overnight on LB+ampicillin plates at 37°C. Colonies were then cultured in 200 µL of LB supplemented with ampicillin in 96-well plates (Sarstedt) and incubated for 8h at 37°C with shaking (200 RPM). For staining, the supernatant was discarded, and the biofilm matrix attached to the wall of wells was gently washed twice with sterile MilliQ water. Then, 300 µl of 0.1% crystal violet stain was added to each well and the plate was incubated for 30 min at room temperature without shaking. After the incubation, the dye was gently discarded, and wells were washed two times with MilliQ water. For dissolving the stained biofilm matrix, 400 µL of 20% acetic acid were added to each well and incubated for 30 min at room temperature without shaking. At the end, 200 µL were transferred to a new 96-well plate (Corning) and the optical density at 600nm was measured using the plate reader. For representative images, the single colonies were grown for 8h in 2.5 mL of LB+ampicillin at 37°C with shaking (200 RPM) in plastic culture tubes (Sarstedt). As above, the supernatant was discarded and the biofilm matrix was gently washed two times with MilliQ water and stained with 5 mL of 0.1% crystal violet for 30 min at room temperature. After two washes with MilliQ water, images were taken.

##### Analysis of fecal Lipocalin-2 levels by ELISA

Inflammation levels were assessed by measuring fecal Lipocalin-2 (LCN-2) using the Mouse Lipocalin-2/NGAL DuoSet kit, as previously described^37^. Briefly, 96-well ELISA plates were coated with capture antibody and incubated overnight at 4°C. The plates were then washed three times with the washing solution PBS-Tween20 (0.05%) and blocked with 1% BSA for 3 h at room temperature. After blocking, the plates were washed three times as before. In parallel, frozen stocks of resuspended feces were thawed on ice, and 100 µL of each sample, along with LCN-2 standards, was added to the plates and incubated overnight at 4°C. The following day, the plates were washed three times before adding the detection antibody. The plates were then incubated for 1 h at room temperature followed by three washes. Next, streptavidin-HRP was added to each well and incubated for 1 hour at room temperature in the dark. The plates were then washed three times. Finally, a 1:1 mixture of TMB and H_2_O_2_ was added, and the plates were incubated for 20 min at room temperature before adding 1M H_2_SO_4_ to stop the reaction. Optical density was measured at 450 nm using a microplate reader.

##### Lipopolysaccharide (LPS) profiling by SDS–PAGE and silver staining

The evolved *E. coli* clones and the corresponding ancestral strains were grown in LB+ampicillin (100 μg/mL) for overnight at 37°C with shaking (200 RPM). For sample preparation, 100 µl of each overnight culture was transferred to 1.5 mL microcentrifuge tubes and centrifuged at 9,500 RPM for 3 min. The supernatant was discarded, and the bacterial pellet was resuspended in 250 µL of SDS loading dye (10 mM Tris-HCl, 2% SDS, 10% glycerol, 0.1% bromophenol blue, pH 6.8). Proteinase K (5 µL, 20 mg/mL) was added, and samples were incubated for 2 h at 58°C. Following incubation, samples were boiled for 5 min at 95°C, and spun down (14000 RPM for 30 sec), and then 10 µL of each sample was loaded onto precast 4–15% Mini-PROTEAN TGX gels (Bio-Rad) and separated by SDS–PAGE in 1× SDS running buffer at 200 V for 30 min. After electrophoresis, the gels were washed twice in ddH₂O for 5 min and fixed overnight in a solution containing 30% ethanol and 10% acetic acid. For silver staining, the gels were incubated for 5 min in an oxidizing solution (2.85 % w/v periodic acid), washed with ddH₂O, and then incubated in Pierce™ Silver Stain solution for up to 45 min with gentle shaking. After two brief washes in ddH₂O, gels were developed using the developer solution until distinct LPS banding patterns became visible. The reaction was stopped by adding 5% acetic acid, and the gels were incubated for an additional 10 min prior to imaging.

##### Phage Isolation

Capsule-dependent phages φYodit and φQuestor (Table S1) were isolated in this study. Pooled wastewater inlet (24 h) from the ARA Basel wastewater treatment plant was used to isolate phages targeting EcST73 and EcST69. Briefly, 50 mL of wastewater was filtered using a 0.2 μm filter, ZnCl_2_ was added to a concentration of 20 mM, and incubated for 1h at 37°C. Precipitates were pelleted by centrifugation and resuspended in 1 mL of LB. Subsequently, 300 μL of the suspension was added directly to 10 mL of warm top agar (LB + 0.5% agar, 50°C) containing EcST73 or EcST69, poured onto a 12 × 12 cm square LB agar plate and incubated overnight at 37°C. The following day, isolated plaques were resuspended in 200 μL of LB and 10x dilutions were spotted on a new lawn of EcST73 or EcST69, and this was repeated twice to ensure clonal phage isolation.

##### Assessment of capsule presence by phage mini-plaque assay and colony morphology on salt-free LB plates

To evaluate the presence or absence of a capsule by susceptibility to capsule-specific phages, plaque assays were performed in a 24-well microplate format as described previously^35^. Briefly, all tested strains, including the wild-type and synthetic capsule-less (Δ*kps)* mutant, were initially grown overnight on LB+ampicillin agar plates at 37°C. Single colonies were picked and inoculated into 50 µL LB in the wells of a 96-well microplate and incubated for 2 hours at 37°C with shaking at 200 RPM. Cultures were then mixed with 250 µL of warm Top Agar and poured in the wells of 24-well microplates, containing 500 µL of 1.5% LB agar. After solidification, 20-50 plaque forming units (PFUs) of either phage φYodit or φQuestor were applied on the Top Agar surface to target EcST73 or EcST69, respectively. Following drying, plates were incubated overnight at 37°C, and the presence or absence of plaques was assessed.

Capsule absence was also assessed from colony morphology on salt-free LB plates, as previously described^37^. Briefly, the evolved clones, together with the wild-type and Δ*kps* control strains, were first grown in LB+ampicillin at 37°C from the cryo-stocks. After overnight growth, the cultures were serially diluted in PBS and 5 μL aliquots were spotted onto salt-free LB plates. Opaque and translucent colonies allowed for distinguishing the presence and absence of capsule, respectively, for each strain.

##### *In silico* prediction of IS elements, prophages, *fimH* and O-antigen typing

To predict mobile genetic elements, namely insertion sequence (IS) elements, present in our *E. coli* strains, all genomes were analyzed using the MobileElementFinder tool^38^ (version 1.03). Prophages were predicted using PHASTEST (version 3.0)^39^. The *fimH* typing was performed using FimTyper (version 1.0)^40^, and serotypeFinder (version 2.0) was used for O-antigen typing^41^.

##### Generation of 3D human urothelial model

Human urothelial tissue models were generated as previously described^42^, with a slight modification. Briefly, cells were seeded onto 6.5 mm, 0.4 µm pore polycarbonate filter membranes in plastic inserts standing in 24-well Transwell plates in CnT-Prime medium. After proliferation, they were exposed to barrier medium (CnT-Prime-3D medium, CELLnTEC) from the basal chamber and human pooled urine from the apical chamber.

##### Urothelial tissue infection and attachment assays

Before infection, all tested strains were maintained statically overnight at 37°C from frozen stocks and then subcultured in fresh LB medium and grown for 48h statically at 37°C. After incubation, bacteria were quantified using QUANTOM Tx Microbial Cell Counter (Logos), according to the manufacturer’s instructions. For infection, bacteria were inoculated in the apical chamber at 10^5^ bacteria/mL in urine and incubated at 37°C, 5% CO_2_. After infection for 2h, the lumen content was collected and tissues washed twice with PBS and incubated with 0.5% Triton X-100 for 20 min. Tissues were then lysed by mechanical disruption, scrapping the surface of the membrane, and collected. Apical milieu and lysates were serially diluted and spread onto square LB agar plates for CFU counting after overnight incubation at 37°C.

##### Assessment of infection bottlenecks *in vivo*

To assess infection bottlenecks *in vivo*, mice were orally gavaged with defined mixtures of untagged wild-type and WISH-tagged *E. coli* strains. WISH-tagged strains were generated based on the previously described wild-type isogenic standardized hybrid (WISH)-tag system^43^, in which the tagged strain carries a neutral chromosomal DNA barcode along with a kanamycin resistance marker, enabling tracking of tagged bacterial populations during infection. For EcST131 I, four WISH-tagged strains, each carrying a unique DNA barcode, were mixed with an untagged parental strain to assess the bottleneck dynamics across a range of tagged inoculum frequencies. In SPF mice, tagged strains were introduced at 10-, 100-, 1000-, and 10000-fold dilution relative to the untagged strain. In LCM and antibiotic-treated LCM mice, tagged strains were introduced at 1000-, 10000-, 100000-, and 1000000-fold dilution relative to the untagged strain. Since all tagged strains carried the same kanamycin-resistance marker, a culture-based quantification on kanamycin-containing plates measured the total tagged population, making these experiments functionally comparable to experiments performed with a single tagged strain. For EcST73, a single WISH-tagged derivative was mixed with the corresponding untagged parental strain at 1:10 and 1:1000 tagged-to-untagged ratios in SPF and LCM mice, respectively. Accordingly, for comparative analyses between EcST131 I and EcST73, EcST131 I datasets were considered equivalent to overall tagged-to-untagged ratios of approximately 1:10 in SPF mice and 1:1000 in LCM mice.

##### Modelling *E. coli* populations growing in the gut of mice

We assume that *E. coli* grow in the gut of mice until they reach carrying capacity, and that periodically, a flushing event occurs, where a large fraction of the bacteria is lost, leading to periodic bottleneck events. We model this process by an alternation of the following two elementary phases:

1. Local growth: Bacteria grow from the bottleneck state, whose average size is denoted by *B*, until they reach carrying capacity *K*.
2. Sampling (flushing): individuals are sampled from the grown population of size *K* using a binomial law, in such a way that we obtain a new bottleneck state where the average size of the population is again *B*. Specifically, individuals are sampled using the following binomial distribution: ℬ(*K*, *B*/*K*).

On average, *B* bacteria are sampled to form the new bottleneck state. The variance of the bottleneck size is given by *B*(1 − *B*/*K*). In the practically relevant case where *K* ≫ *B*, the binomial law can be approximated by a Poisson law PP(*B*), and the variance of the bottleneck size by *B*.

##### Bottleneck size estimation

To estimate the bottleneck size *B* from each of the experiments shown in **Figure S18**, we focus on the extinction times of the tagged strain in each mouse (i.e., the times where the tagged strain can no longer be detected). Briefly, we express the likelihood of observing such extinction times under a neutral Wright-Fisher model with population size *B*. Next, we maximize this likelihood, to obtain the value of *B* that best explains the data.

As detailed above, the population undergoes cycles of growth and sampling (flushing) events, and the latter can be modeled by binomial sampling. For mathematical tractability, we will however calculate the bottleneck size under the approximation that it is fixed, meaning that at each bottleneck event, *exactly B* bacteria are sampled (instead of a number whose average is *B* but that can fluctuate), and we will also assume *K* ≫ *B* and neglect the finite size of the grown population (i.e., we make the approximation *K* → ∞). This allows us to approximate our model by a Wright-Fisher model with population size *B*. Tagged strains are assumed to be neutral, in the sense that tags are assumed not to impact fitness of *E. coli* bacteria. Hence, when a tagged strain is introduced at proportion (or dilution factor) *x*, we can consider it as a neutral mutant drifting in the population with initial fraction *x*. The extinction of tagged strains then corresponds to the extinction of a neutral mutant lineage under the Wright-Fisher model, which is a classical problem in population genetics.

Under Kimura’s diffusion approximation, the cumulative distribution of the extinction time *T*, expressed in number of bottleneck events, of a neutral mutant lineage in the Wright-Fisher model for a haploid population can be expressed as^44,45^:

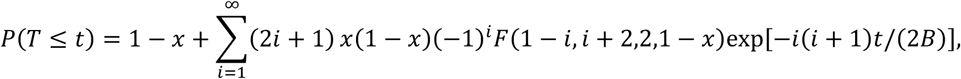

where *FF* denotes the (ordinary) hypergeometric function. This quantity represents the probability that the tagged strain gets extinct by time *t* (i.e. that its extinction time *T* is smaller than or equal to *t*). It depends on the initial proportion *x* of the tagged strain, which is known, and on the bottleneck size *B*, which we are aiming to estimate from the data.

In each experiment, we consider the data as a list of the number *n_t_* of mice where the tagged strain was lost at time *t*. We also take into account the fact that some mice do not lose their tagged strains before the end of the experiment, which occurs at a time denoted by *τ*. The number of such mice is denoted by *n_R_* (for “remaining”). Since the complete renewal of the cecum content (where most of the growing E. coli population is located) takes about 16h^46^, we assume that there is one bottleneck per day. Hence, times in numbers of days are interpreted as times in numbers of bottleneck events. Considering each mouse as independent, the likelihood of the data can then be expressed as:

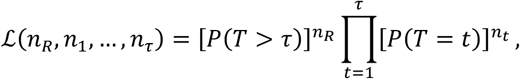

where the probability that extinction happens at bottleneck *t* is obtained from our above expression of

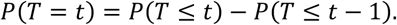

In practice, because it is mathematically equivalent but numerically more convenient, we maximize the log-likelihood log ℒ(*n_R_*, *n*_1_, …, *n_τ_*), which reads:

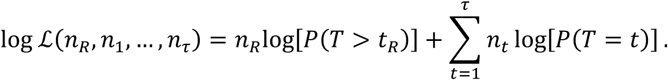

For each experiment reported in **Figure S18**, we determine the value of *B* that maximizes this log-likelihood.

We obtain bottleneck size estimates of *B* = 3.7 × 10^5^ for LCM mice colonized by strain EcST131 I (see **Figure S19A**, where the data from **Figure S18E** was used), *B* = 8.9 SPF mice colonized by strain EcST131 I (see **Figure S19B**, where the data from **Figure S18A** was used) and *B* = 39.6 for SPF mice colonized by strain EcST73 (see **Figure S19C**, where the data from **Figure S18B** was used). Note that cases where the tagged strain is not lost in any mouse do not allow maximum-likelihood estimation of *B* (only providing lower bounds). This is the case for antibiotic-treated LCM mice, with both initial fractions of tagged strains considered (see **Figures S18D and F**). In these cases, we can only say that the bottleneck size is greater than in LCM mice.

In **Figure 3E**, we report the measured carrying capacity *K* versus the estimated bottleneck *B* in three different cases. We observe a nonlinear scaling of *K* versus *B*.

##### Extinction probability due to stochastic sampling at each bottleneck

Extinctions of the bacterial population can happen at each flushing phase, if all bacteria are flushed out of the system. This is a stochastic effect which is a direct consequence of the randomness intrinsic to a flushing event. In our model, where individuals are sampled using a binomial law ℬ(*K*, *B*/*K*), the probability that 0 bacteria are sampled to form the new bottleneck state reads:

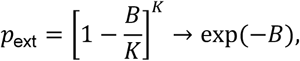

where we took the limit where *K* → ∞ in the last step. Note that using the Poisson approximation of the binomial law directly leads to the same result. This expression was used to plot **Figure 3F**.

Since extinction of the population happens with probability *p*_ext_ at each bottleneck, the number of bottlenecks needed to obtain extinction of the population follows a geometric distribution with mean ⟨*t*⟩ = 1/*p*_ext_ = exp(*B*). For *B* = 8.9, which is our estimate of the bottleneck size for SPF EcST131 I, this yields ⟨*t*⟩ = 7.3 × 10^3^ bottlenecks (i.e., days). While this remains longer than the evolution experiments described here, *B* can in fact vary in time. We note that for *B* = 5, we have ⟨*t*⟩ = 148 bottlenecks (i.e., days). Hence, our model predicts that extinction events are frequent on the timescale of the evolution experiments reported here, if bottlenecks are just a little bit smaller than those estimated for SPF EcST131 I.

---

**Figure S1.**
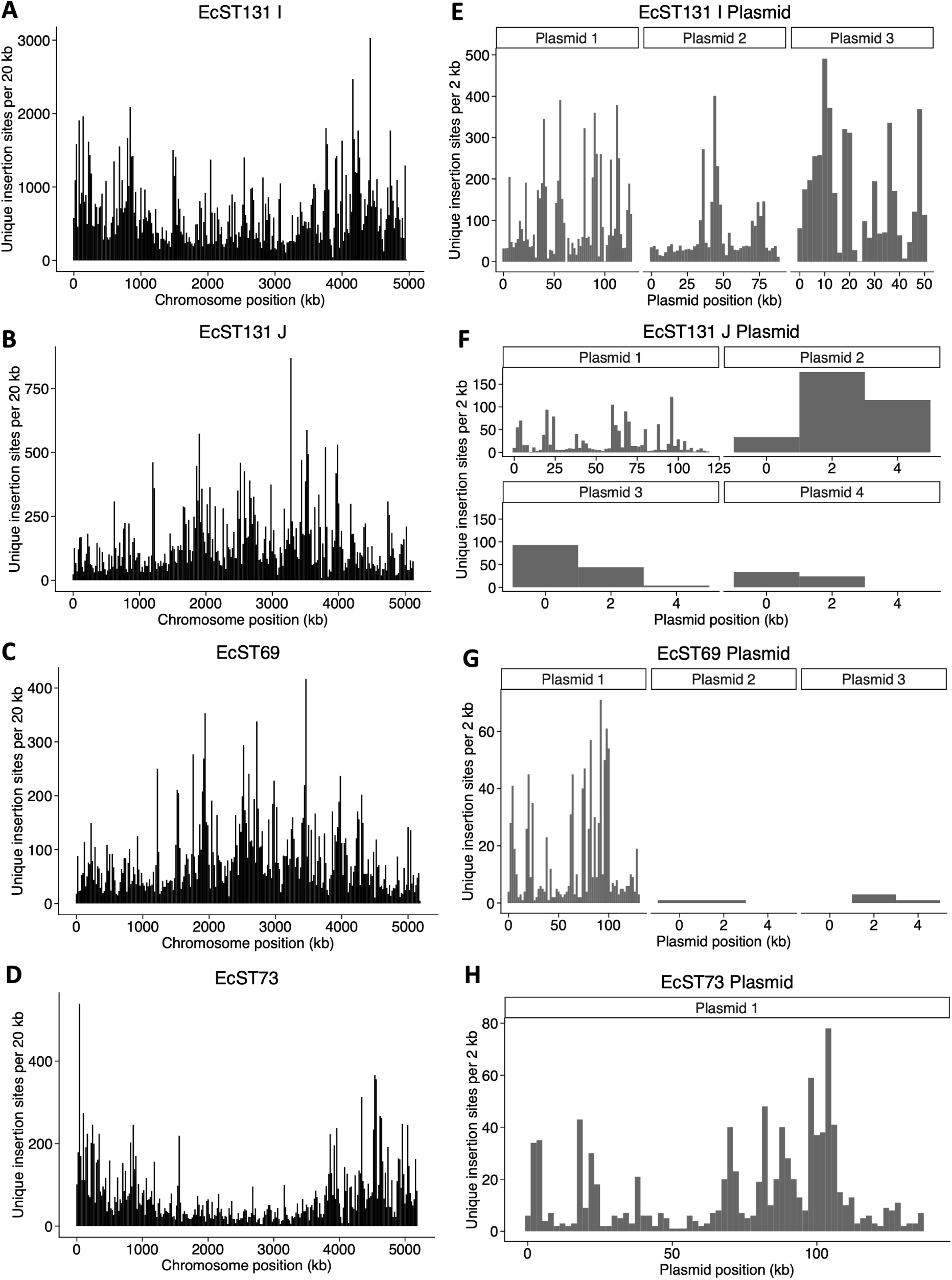
Genome-wide distribution of unique RB-TnSeq insertion sites on chromosomes (A-D) and plasmids (E-H). Bars represent the number of unique insertions per 20 kb across the chromosomes **(A-D)** and per 2 kb across each plasmid **(E-H)**. The genome coverage was 92.45%, 76.65%, 67.79%, and 63.41% for strain EcST131 I, EcST131 J, EcST69, and EcST73, respectively. The median number of insertions per genomic feature ranged from 3 to 21 (see Table S2).

**Table S2:** Summary of RB-TnSeq insertion data for each strain.

| **Strain** | **Number of insertions**<br>(chromosome + plasmids) | **Genomic feature coverage**<br>(number of features with<br>insertion (chromosome +<br>plasmids) / total features | **Median insertions per**<br>**genomic feature**<br>(chromosome+plasmids) |
| --- | --- | --- | --- |
| EcST131 I | 326846 (297110+29736) | 92.45% (4889/5288) | 21 |
| EcST131 J | 53496 (50509 + 2987) | 76.65% (4079/ 5321) | 5 |
| EcST69 | 27372 (26173+ 1199 | 67.79% (3661/5400) | 3 |
| EcST73 | 26128 (24912 + 1216) | 63.41% (3370/5314) | 3 |

Genomic features were defined as all annotated entries in the genome annotation file, including coding sequences (CDS), tRNA, rRNA, and misc\_RNA. All features were included in the transposon insertion mapping analysis.

**Figure S2.**
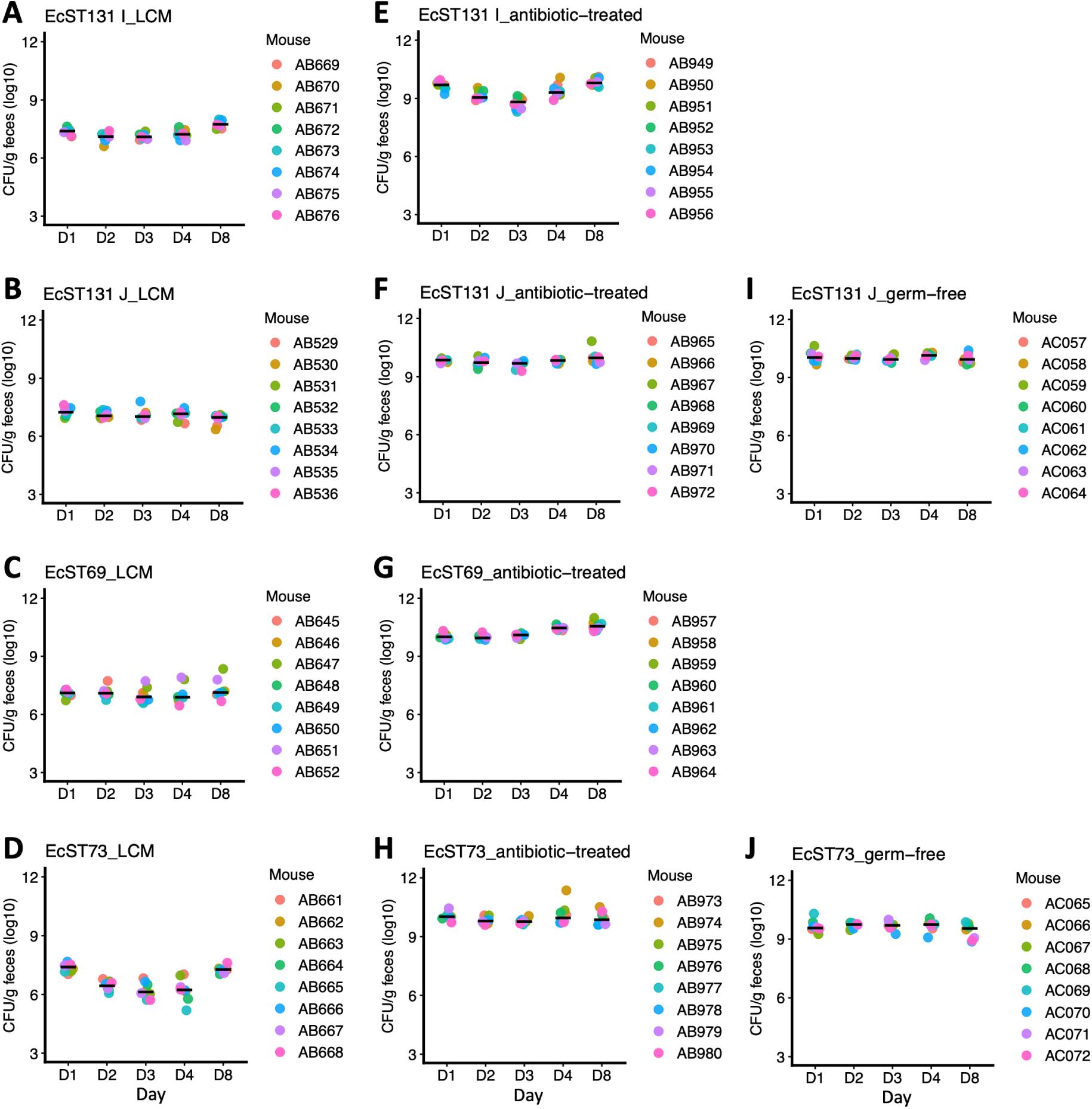
Total bacterial load in feces of LCM (A-D), antibiotic-treated LCM (E-H), and germ-free (I and J) mice orally inoculated with the mutant library. Each data point represents one mouse, and the horizontal line shows the median.

**Figure S3.**
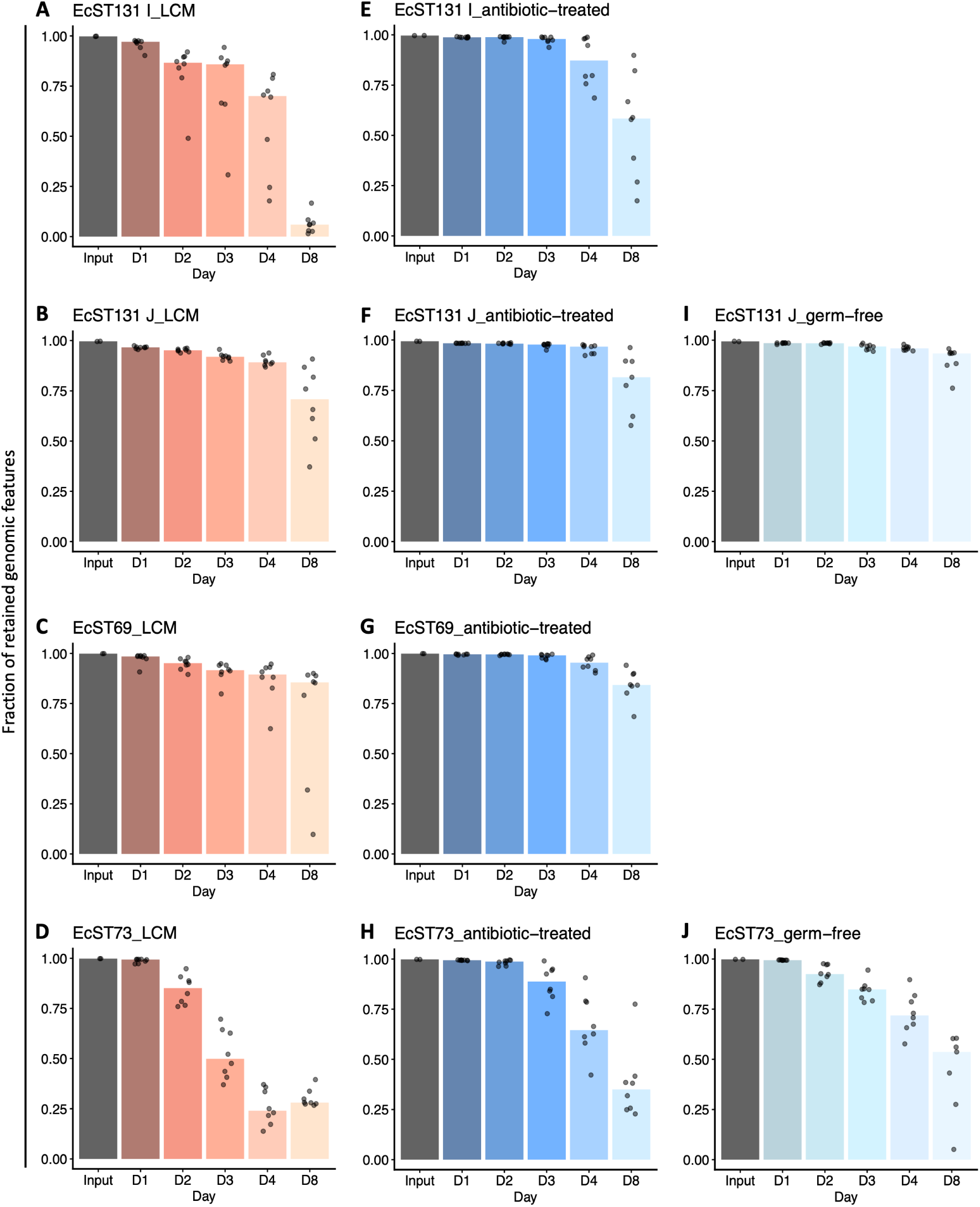
Changes in retention of genomic features over time across strains and microbiota conditions. Fraction of retained genomic features detected by library sequencing before (input) and after infection of LCM **(A-D)**, antibiotic-treated LCM **(E-H)**, and germ-free mice **(I and J)**. Genomic features refer collectively to genes, non-coding RNA, and intergenic regions represented in the RB-TnSeq libraries. The fraction retained was calculated as the number of features detected in a given sample divided by the total number of unique features detected across both inoculum replicates. The retention of genomic features remains largely stable for early time points (D1/D2), suggesting no detectable bottleneck. Therefore, for downstream analyses, we selected the latest time points that still showed stable retention for each strain and condition, maximizing infection duration while avoiding bottleneck effects: D1 for EcST131 I and EcST73, and D2 for EcST131 J and EcST69. Bars represent the median fraction of retained features, and each point represents an individual mouse sample (*n*=8).

**Figure S4.**
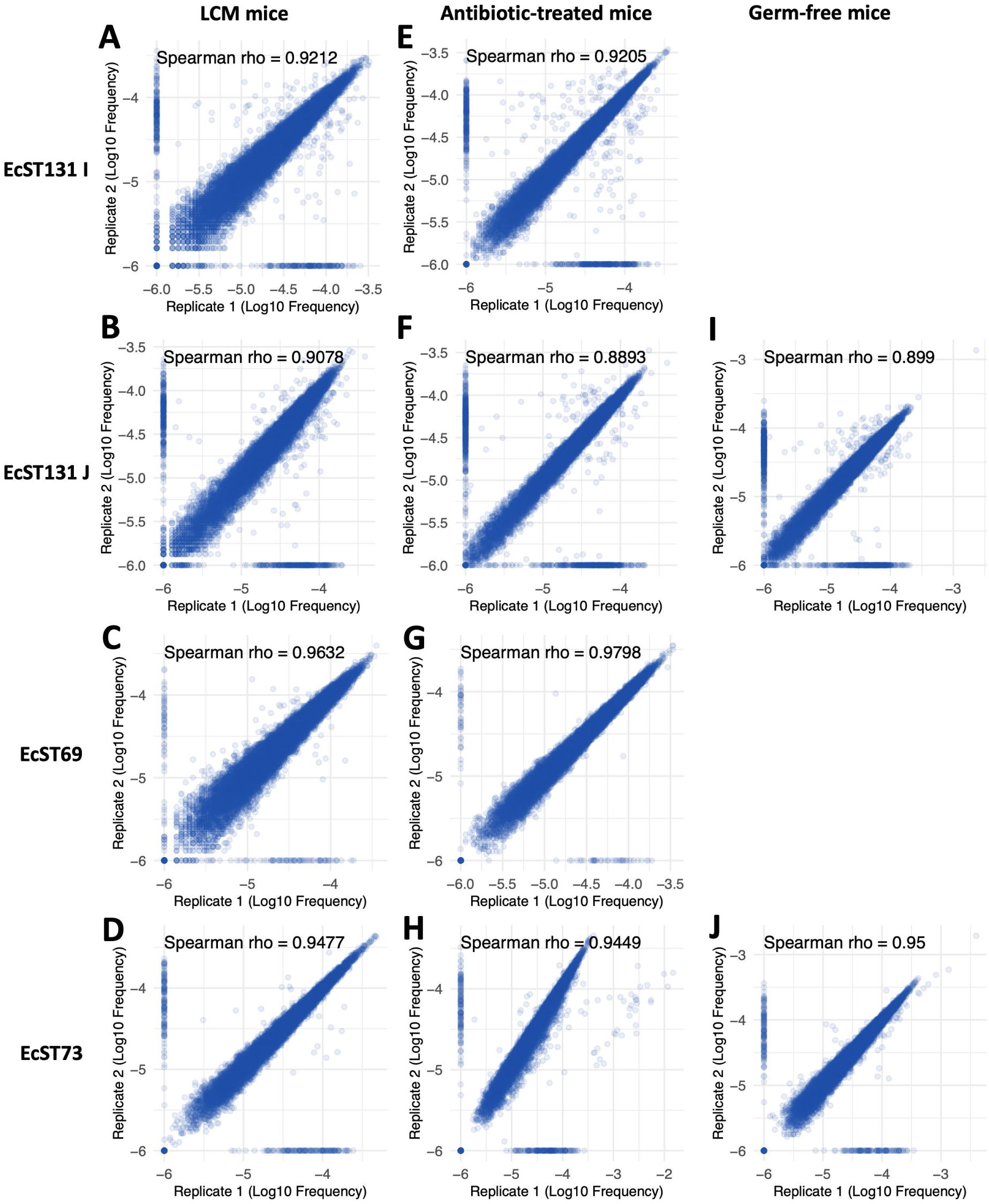
Library preparation and sequencing are highly reproducible. Correlation plot of mutant frequencies between input library replicates. Each dot represents a single mutant detected in the input pool.

**Figure S5.**
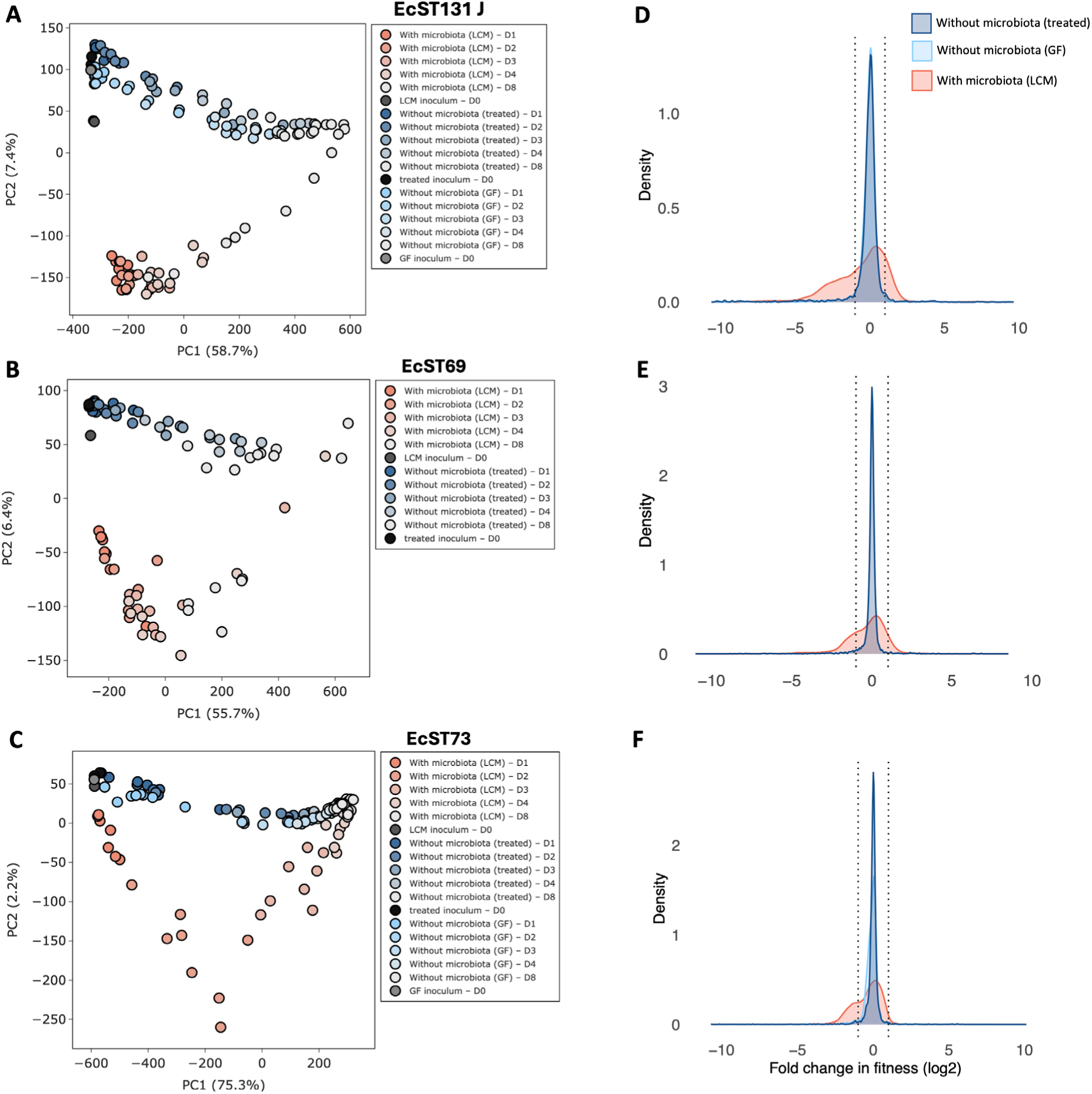
Effect of the gut microbiota on *in vivo* fitness landscape of ESBL *E. coli*. **(A-C)** Principal component analysis of RB-TnSeq profiles across screening conditions for each strain, based on normalized transposon barcode abundances (see Methods). Each point represents an individual mouse sample. **(D-F)** Distribution of fitness effects (DFEs) of genes during bacterial colonization in the presence (red) and absence of microbiota (blue- antibiotic-treated; cyan – germ-free). Histograms are shown after Gaussian kernel density smoothing. Dashed lines indicate neutral fitness window.

**Figure S6.**
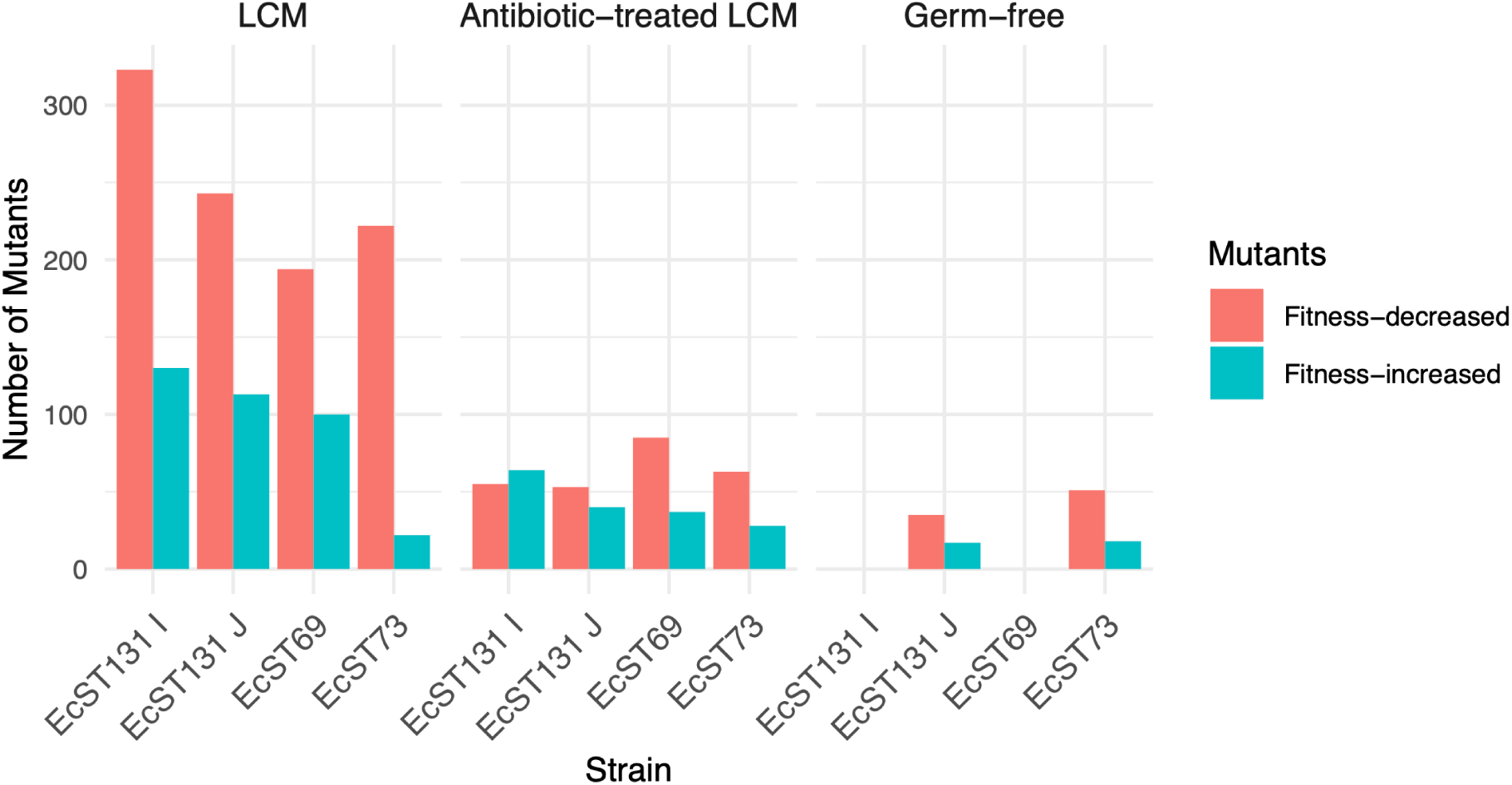
Total number of mutants that showed a significant change in fitness in the RB-TnSeq screen. Bar plots show the total number of mutants identified by RB-TnSeq as significant hits: mutants showing decreased fitness (red) and increased fitness (teal).

**Figure S7.**
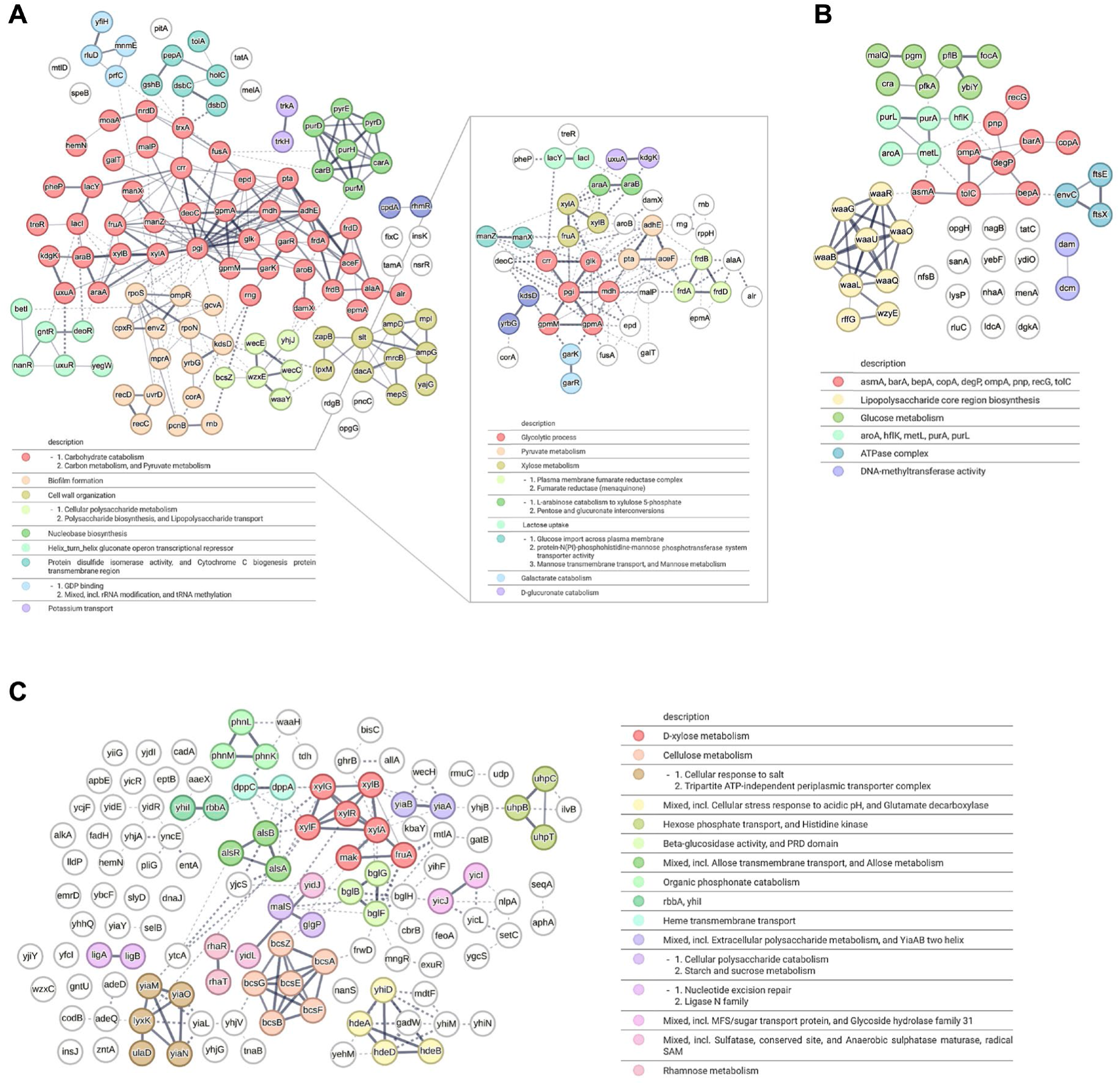
Genes altering *in vivo* fitness. Interaction network for mutants exhibiting a fitness defect (in at least one strain) in antibiotic-treated mice **(A)**, and common in both LCM and antibiotic-treated (and/or in germ-free mice) **(B)**. Panel **(C)** shows mutants exhibiting increased fitness in LCM mice. Protein-protein interactions were obtained with STRING.

**Figure S8.**
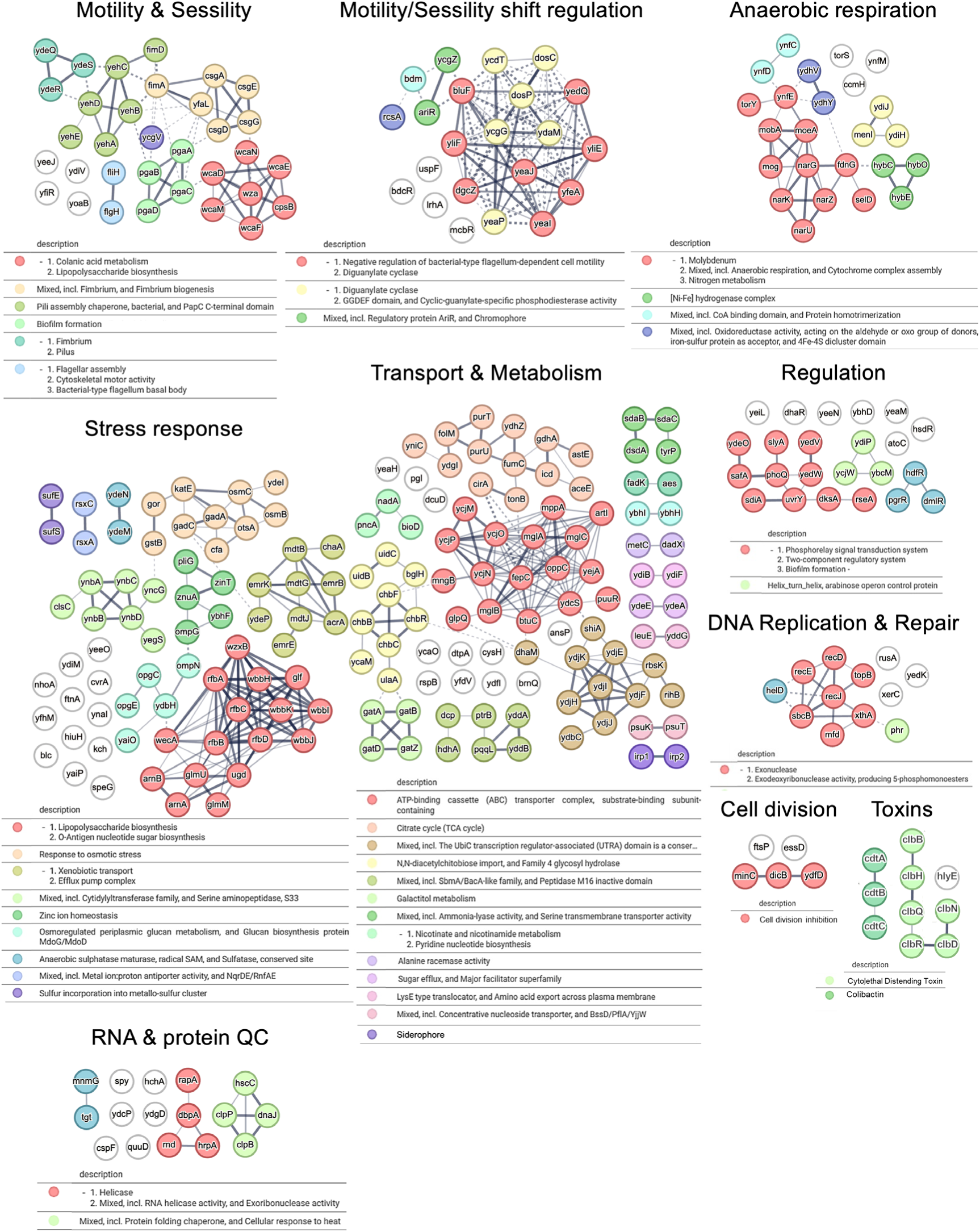
Genes altering *in vivo* fitness in the presence of microbiota. Interaction network for mutants that showed fitness defect, in at least one strain, specifically in the presence of gut microbiota. The protein-protein interactions network was generated using STRING (see Methods).

**Figure S9.**
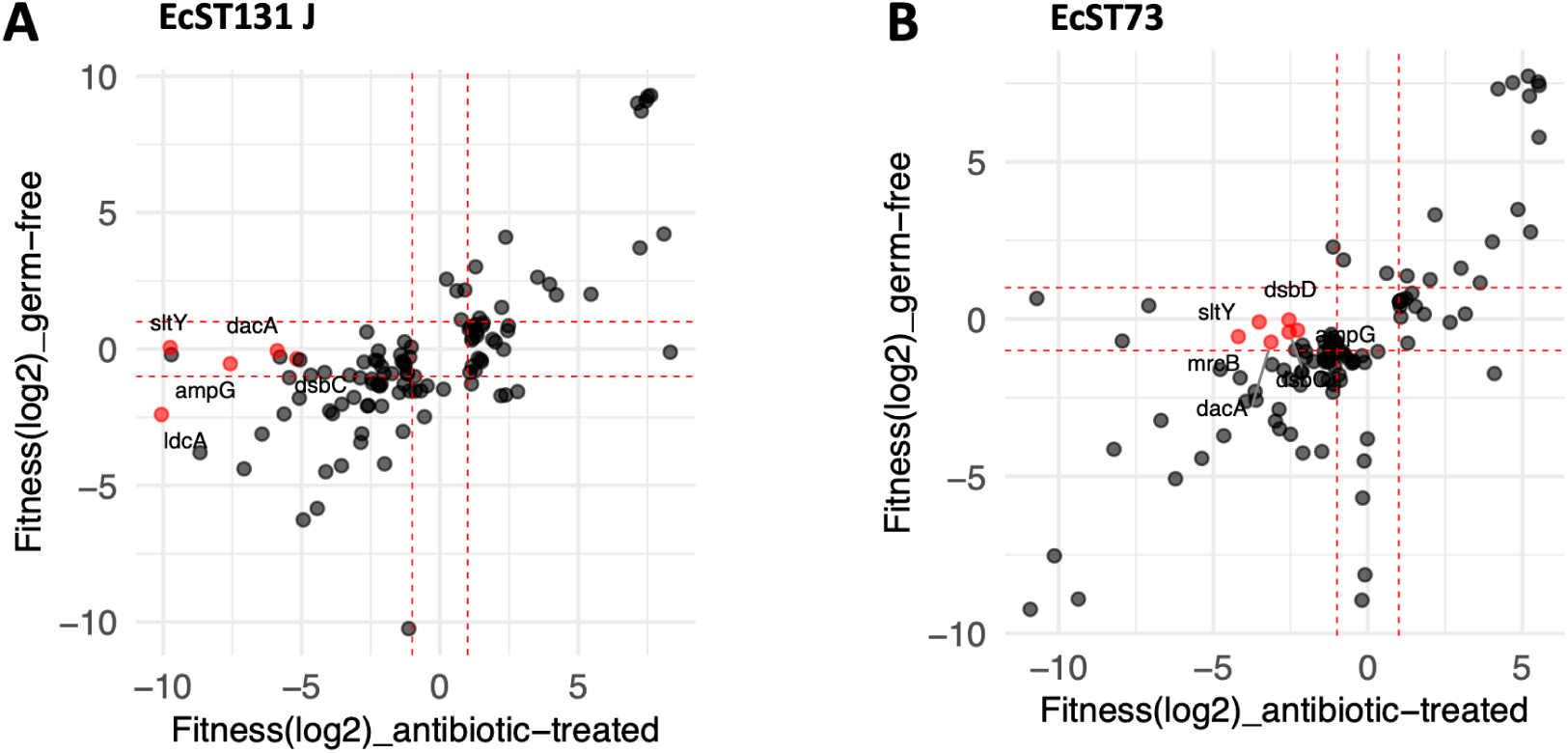
Correlation of fitness scores between antibiotic-treated LCM and germ-free conditions. Panels **(A)** and **(B)** show correlation restricted to significant hits (fitness-decreasing and fitness-increasing genes) of EcST131 J and EcST73, respectively. Each dot represents a single gene. Dotted lines indicate the significance thresholds for fitness effects in each condition. Red dots highlight some fitness-decreasing genes specific to the antibiotic-treated condition and potentially are affected by ampicillin treatment, as these genes are involved in peptidoglycan biosynthesis/recycling or maturation. Spearman’s rank correlation coefficients (*r_S_*) and corresponding *p* values: (A) EcST131 J: *r_s_* = 0.644, *p* = <2.2e-16; and (B) EcST73: *r_s_* = 0.60, *p* = <2.2e-16.

**Figure S10.**
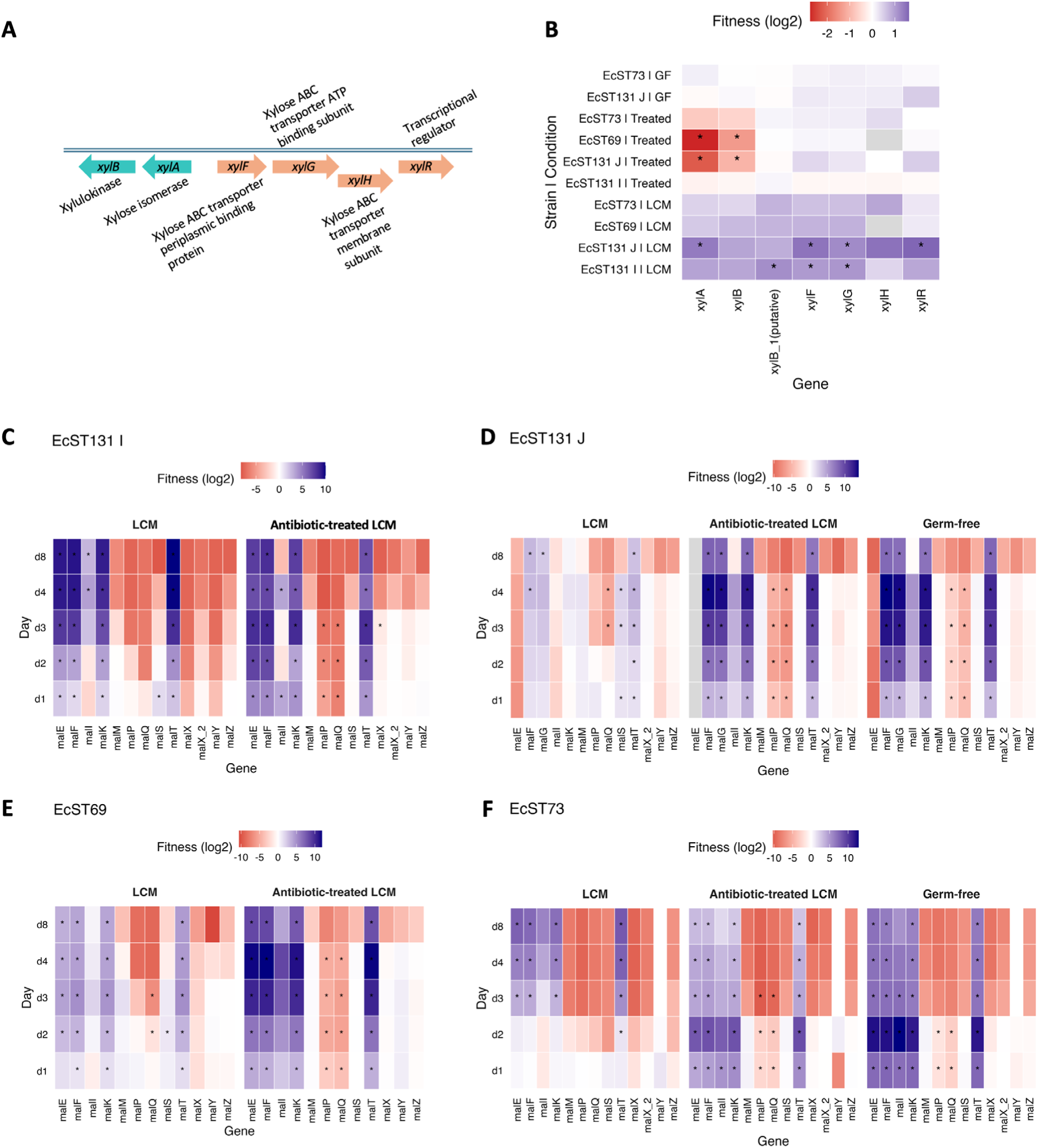
Fitness values of xylose and maltose utilization/uptake mutants. **(A)** Schematic of the xylose utilization and transport operon. Heatmaps show RB-TnSeq-derived fitness scores for xylose **(B)** and maltose uptake- and metabolism-related mutants across strains and conditions **(C-F)**. Color scale indicates fitness score (log_2_ fold change), where positive values represent increased mutant fitness and negative values represent fitness defects. Asterisks denote mutants with statistically significant changes in fitness in the screen.

**Figure S11.**
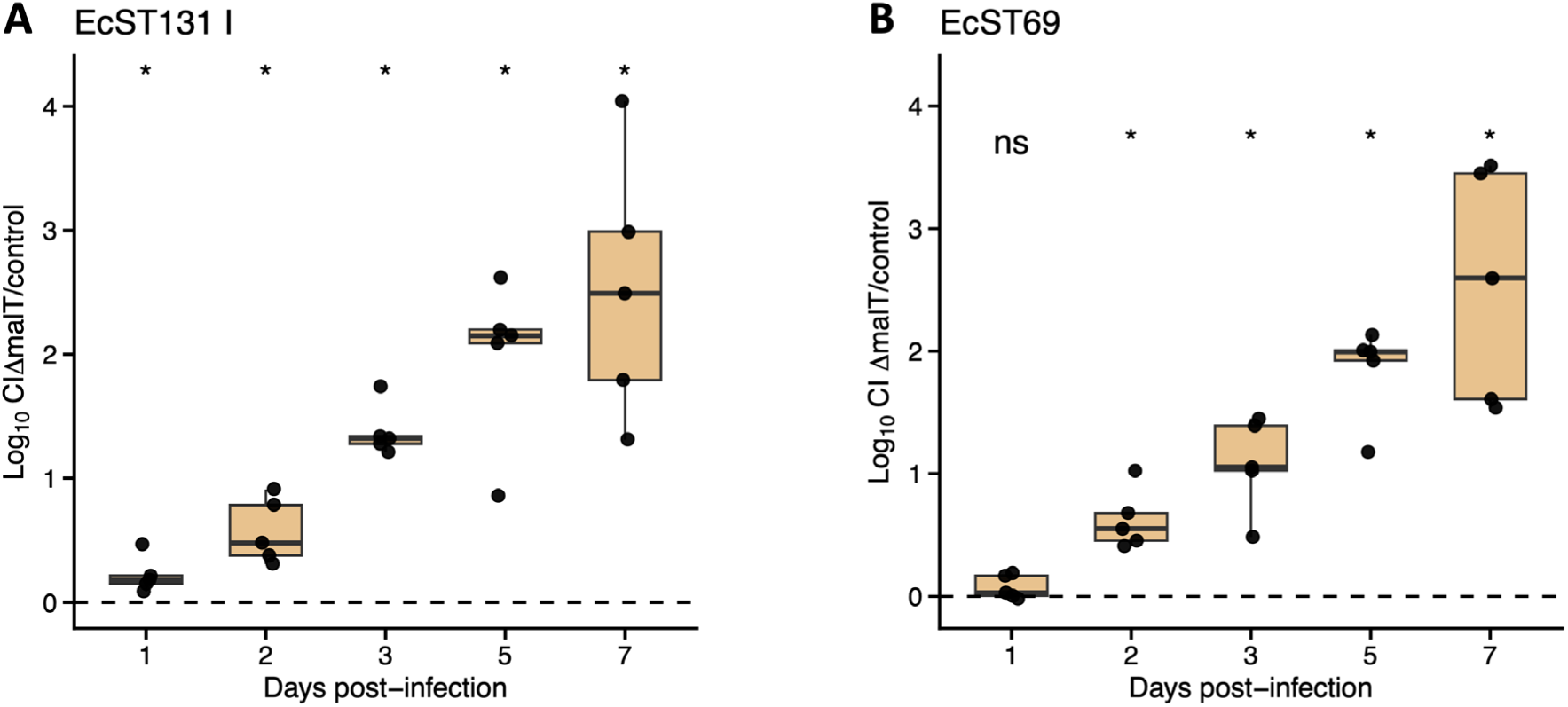
*In vivo* competitive fitness of Δ*malT* mutant. Boxplots showing competitive indices from *in vivo* competition experiments in LCM mice (*n*=5) with **(A)** EcST131 I and **(B)** EcST69. Each point represents an individual mouse. Statistical significance was determined using one-sided Wilcoxon signed-rank test and significance levels are indicated as ns-not significant, \**p* <0.05.

**Figure S12.**
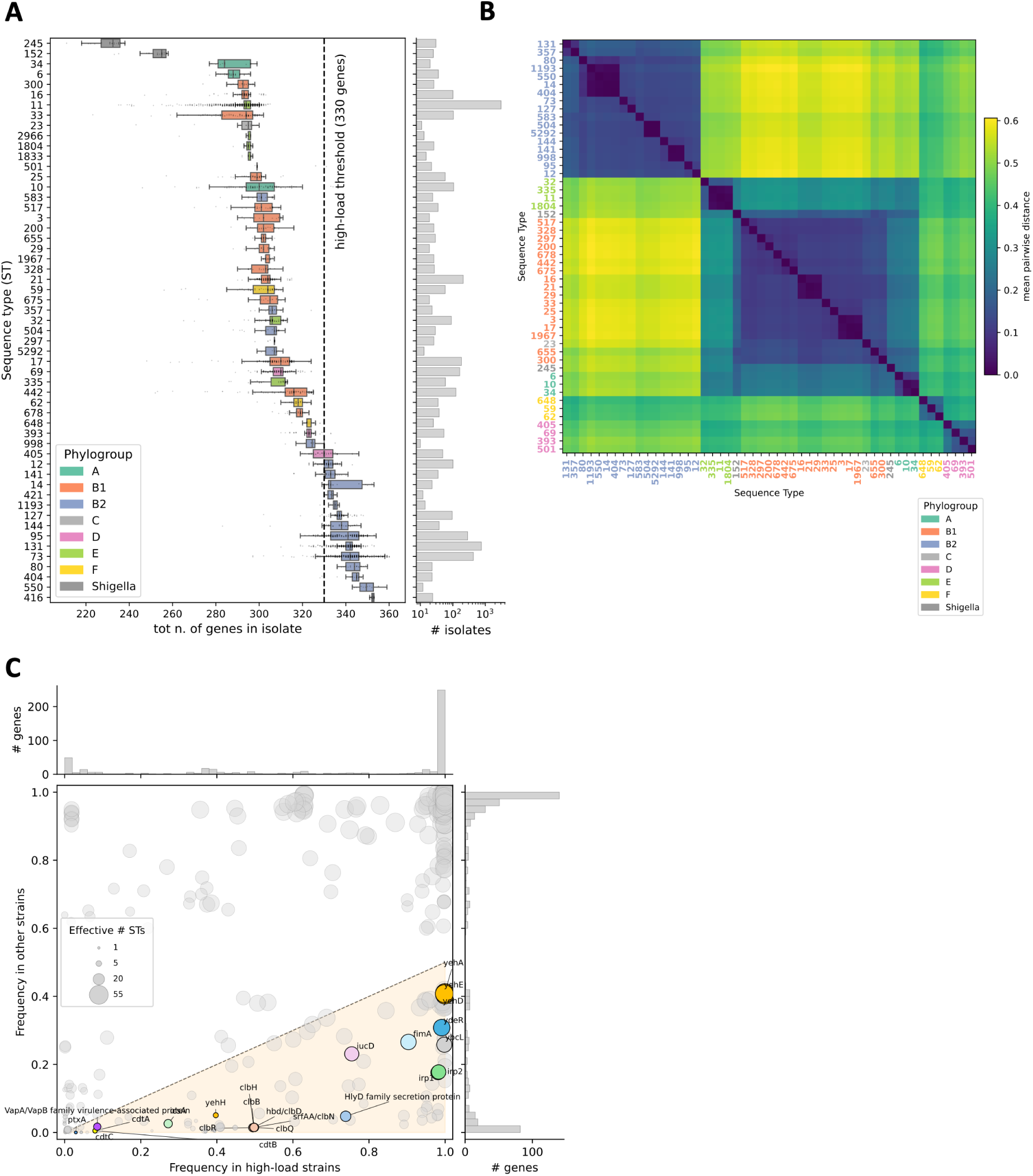
Genes that increase *E. coli* fitness in the presence of microbiota are prevalent in most STs belonging to phylogroup B2. **(A)** Box plots show the distribution of genes that increase fitness specifically in the presence of gut microbiota across genomes, representing different STs and phylogroups (Horesh collection^19^, see Methods). Only STs with ≥10 isolates were included in the analysis. Each box is colored by the phylogroup of the ST. The vertical dashed line marks the high-load threshold of 330 genes (see Methods). **(B)** Average tip-to-tip distance between isolates from different STs in the Horesh core-genome tree, calculated as detailed in methods. ST labels are coloured by phylogroup. **(C)** Per-gene frequency in high-load strains (*n*= 1′878) against frequency in normal-load strains (*n*= 5′634), across 485 genes that increase fitness in the presence of microbiota. Marker size encodes the ST participation ratio, calculated as described in the methods. The orange wedge marks the region of ≥ 2-fold enrichment in high-load strains, freq_high-load_ > 2 ⋅ freq_normal-load_. Selected virulence-associated and pathoadaptive genes are highlighted. Marginal histograms on the top and right show the distribution of each frequency across all 485 genes.

**Figure S13.**
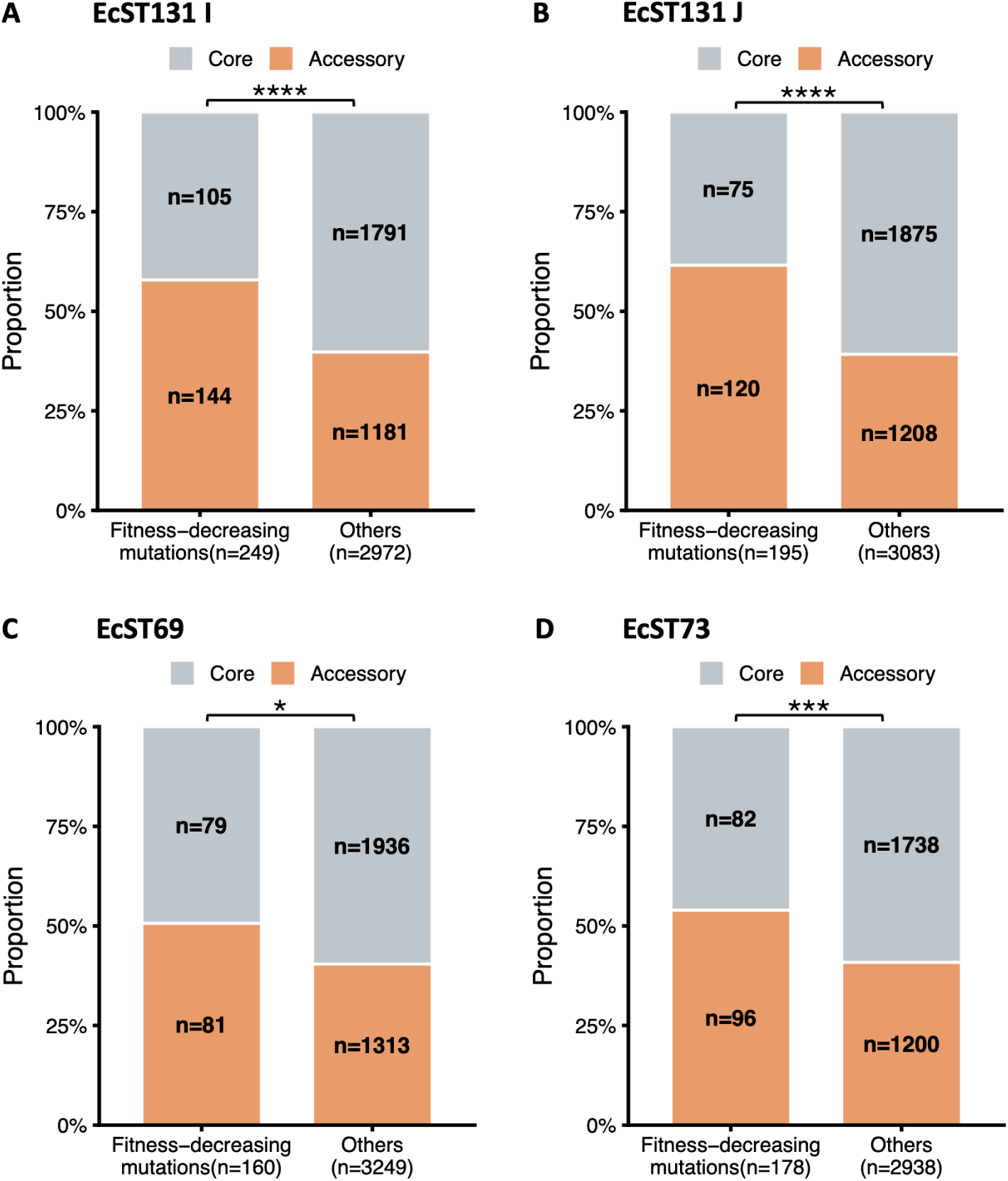
Accessory genes are significantly enriched among genes that increase fitness in the presence of microbiota. Stacked bar plots show the proportion of core and accessory genes among microbiota-specific (LCM) fitness-decreasing mutations and other mutations (i.e., fitness-increasing mutations, neutral mutations, and fitness-decreasing mutations in both microbiota-colonized and microbiota-depleted conditions). Enrichment of accessory genes in fitness-decreasing mutations was assessed using Fisher’s exact test for each strain. \**p* <0.05, \*\*\**p* <0.001, \*\*\*\**p* <0.0001.

**Figure S14.**
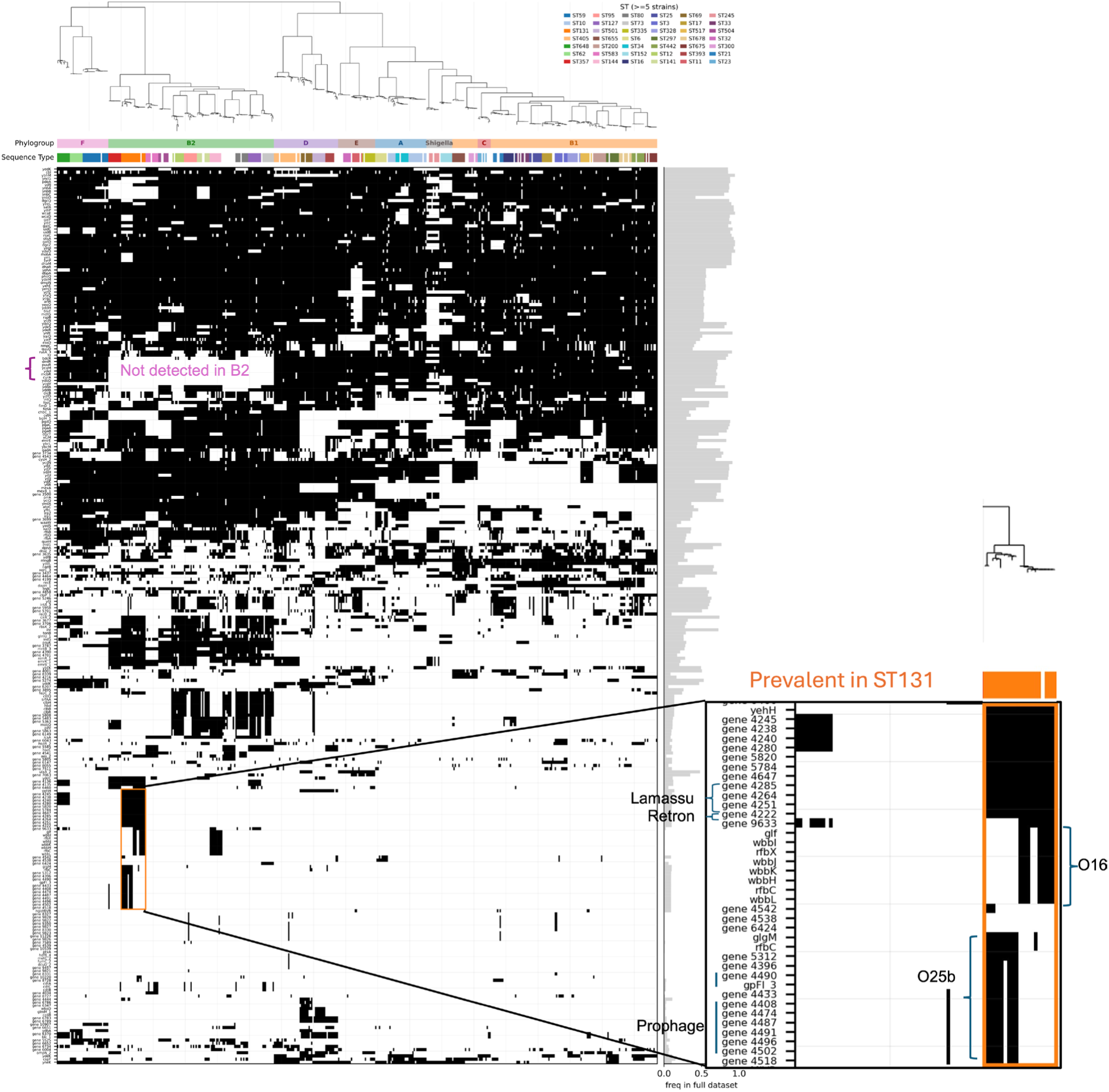
Distribution of genes that increase fitness in the presence of microbiota across diverse *E. coli* isolates. Presence/absence of the 282 genes that increase fitness in the presence of microbiota (the ones with full-dataset frequency < 0.95, to exclude core genes) across the isolates present in the Horesh core-genome tree (on top). STs with ≥ 5 isolates in the tree are coloured according to the legend on the right. Genes are ordered by hierarchical clustering on the full-dataset presence/absence matrix. The right-hand bar plot reports the frequency of each gene in the full dataset. The box highlights genes of interest that are prevalent in ST131.

**Figure S15.**
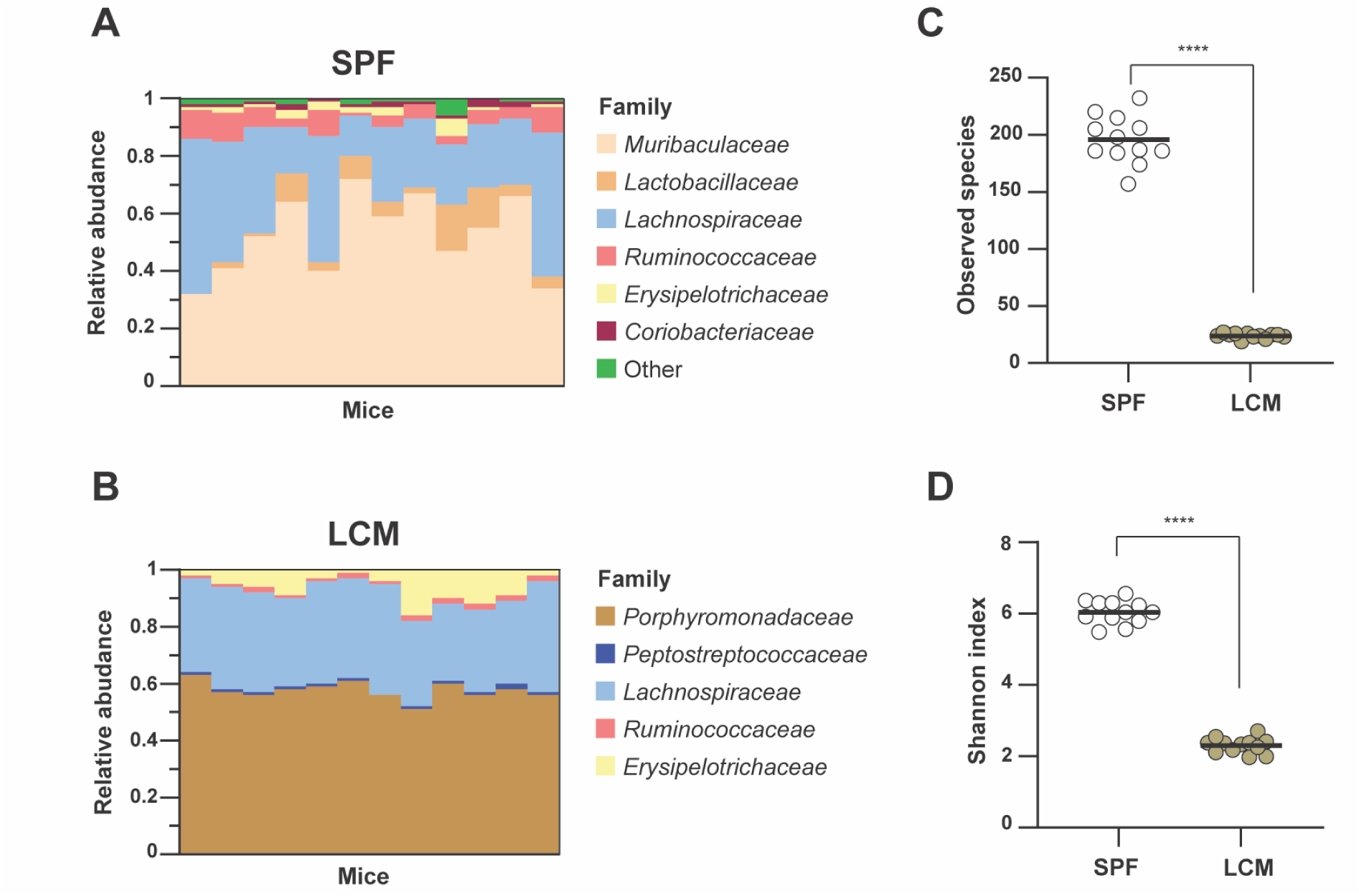
Microbiota analysis highlights the higher complexity of SPF gut microbiota in comparison to LCM mice and the lack of Enterobacteriaceae in these models. Relative abundance of all detected taxonomic families isolated from **(A)** SPF and **(B)** LCM mice feces and subjected to 16S amplicon sequencing. **(C)** Number of observed species for SPF and LCM samples. **(D)** Shannon index for microbiota diversity for SPF and LCM samples. Statistics were performed with Mann-Whitney tests, \*\*\*\**p* <0.0001.

**Figure S16.**
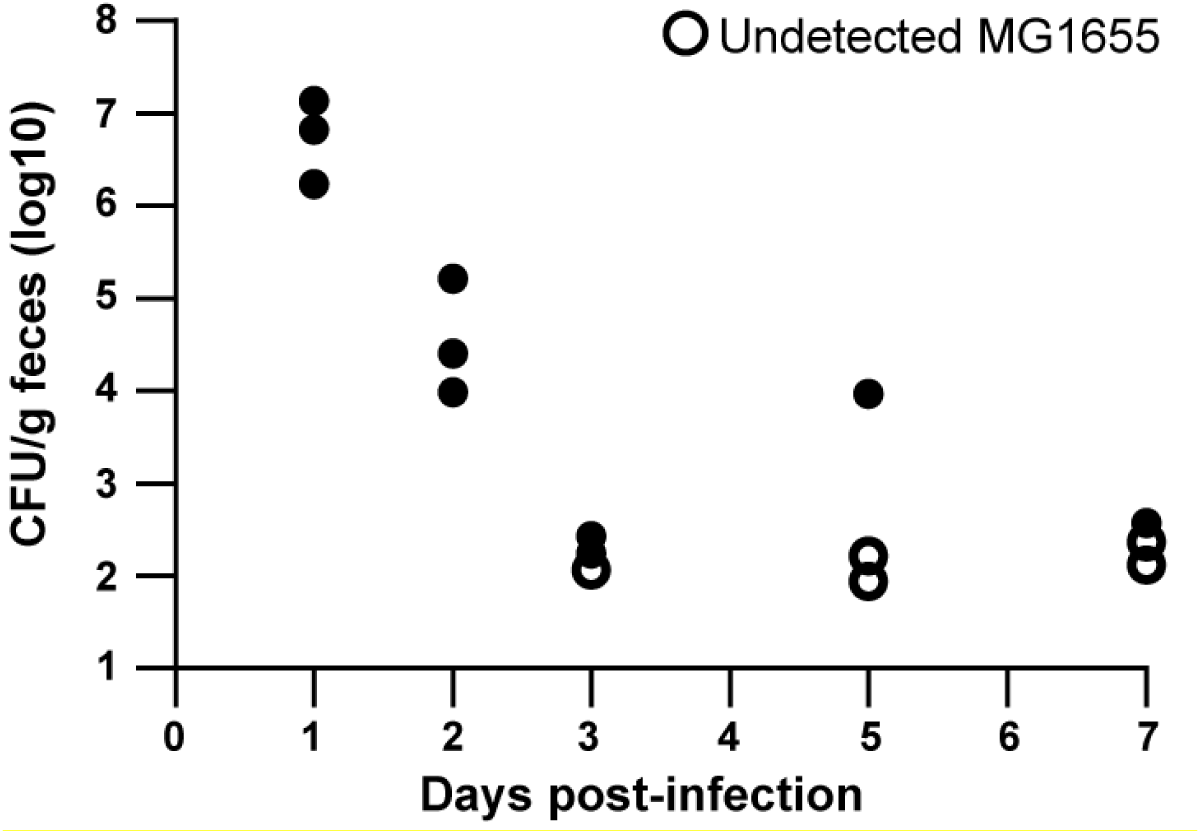
*E. coli* MG1655 laboratory strain failed to stably colonize conventional SPF mice over one week. Colonization levels are shown as CFU per g feces of *E. coli* MG1655 with restored O-antigen. Each point represents an individual mouse (*n*=3), and open circles denote samples in which *E. coli* was below the limit of detection.

**Figure S17.**
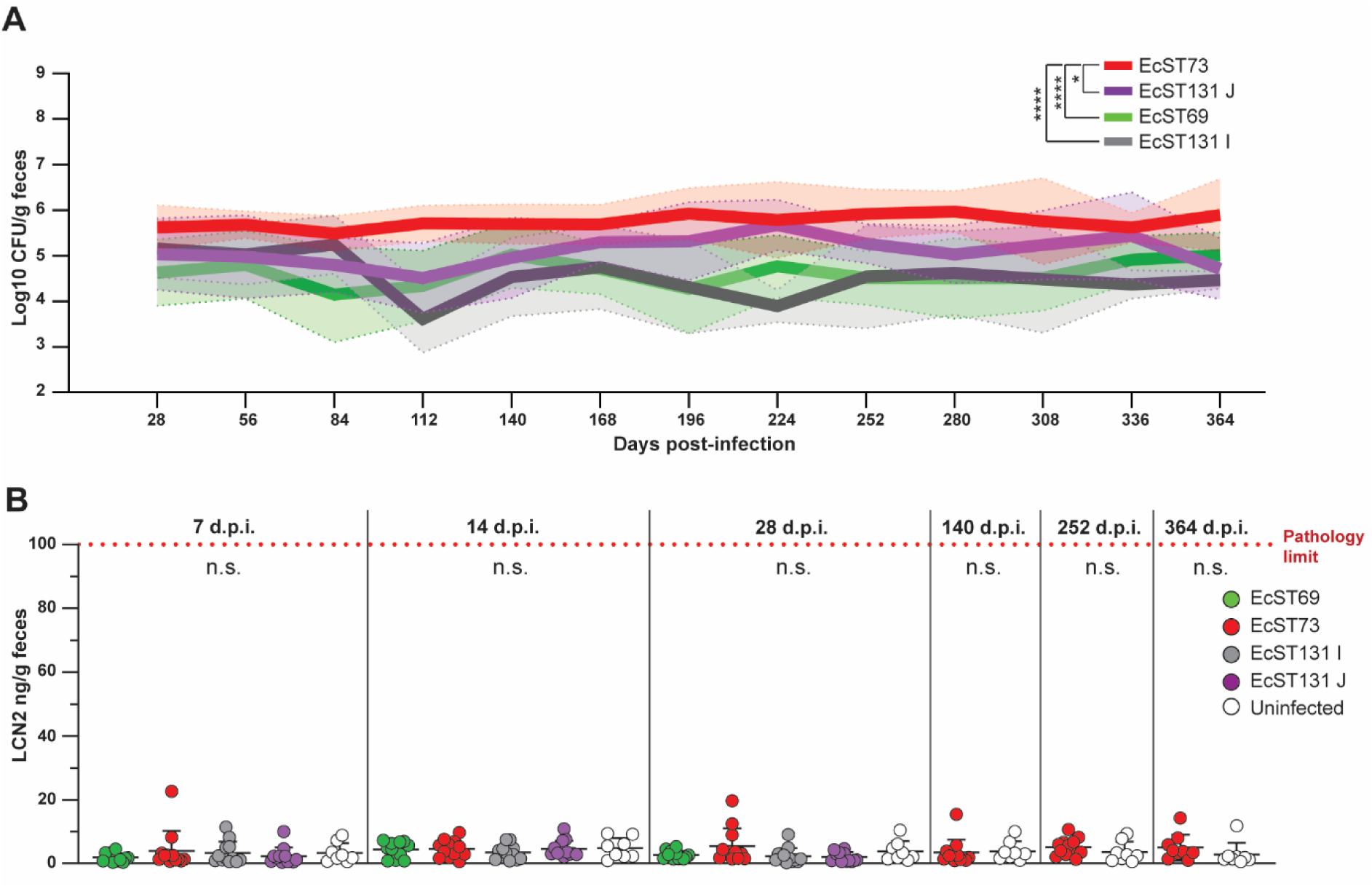
In SPF mice, EcST73 exhibits a higher carrying capacity than the other tested strains without signs of gut inflammation. **(A)** EcST73 reaches higher colonization levels than the other tested strains. Colonization levels (Log10 CFU/g feces) of detectable EcST73 during the evolutionary experiment are shown. Bold lines represent mean colonization levels for each strain, and shaded areas indicate the standard deviation. Statistical analysis was performed using one-way ANOVA of area under the curve (AUC) data, with Dunnett’s multiple comparisons test and EcST73 as the control group; * *p*<0.05, **** *p*<0.0001. **(B)** Colonization of SPF mice by EcST131 I, EcST131 J, EcST69, and EcST73 did not induce gut inflammation. Fecal LCN2 levels (LCN2 per g of feces) were measured in mice colonized with different strains (*n*=12) and in uninfected control mice (*n*=8). Statistical analysis was performed using one-way ANOVA with Dunnett’s multiple comparisons test at each time-point, comparing colonized mice with uninfected control mice. Red dotted-line denotes the minimum pathology limit; n.s. means not significant.

**Figure S18.**
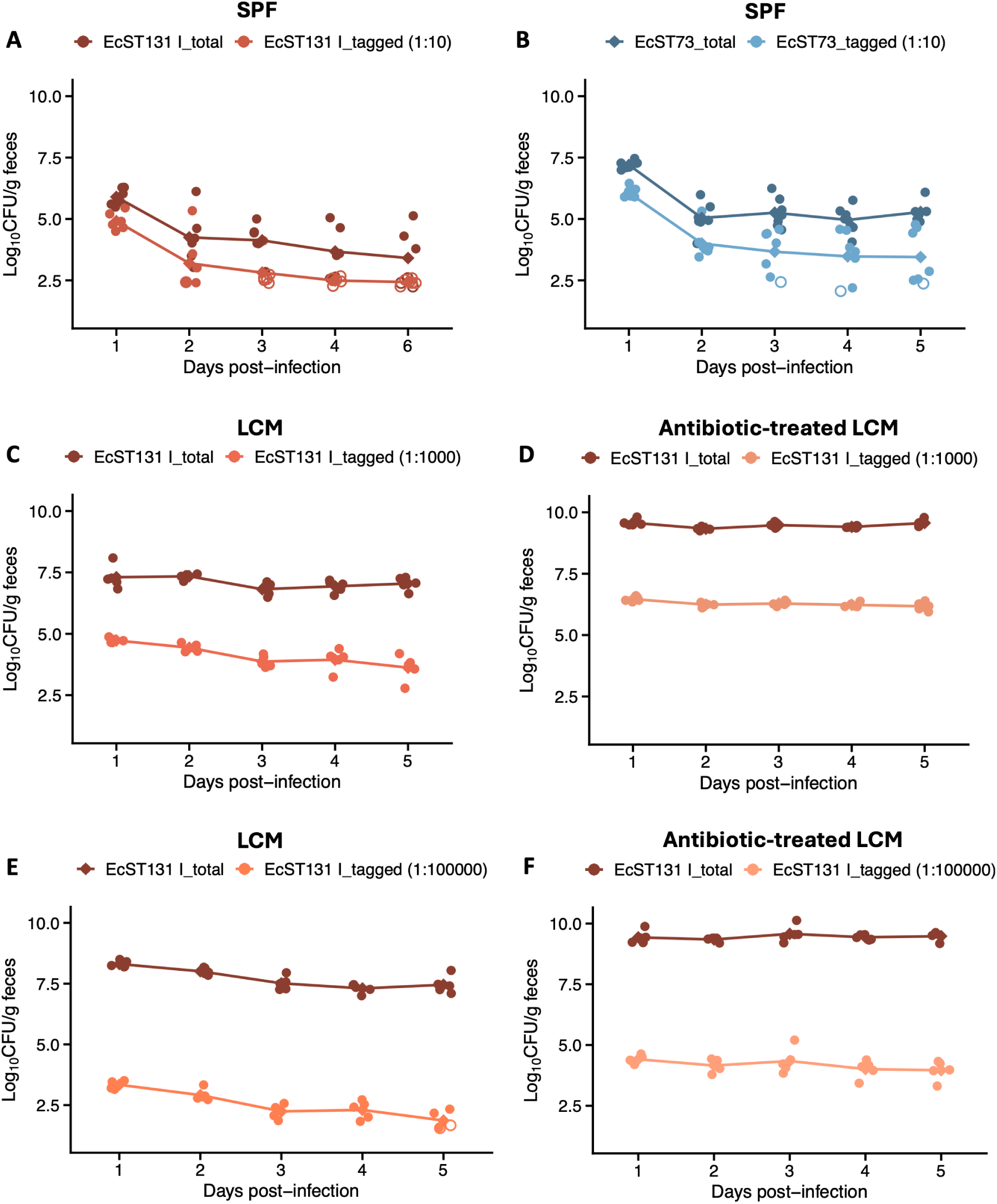
*In vivo* bottleneck analysis. SPF **(A and B)**, LCM **(C and E)**, and antibiotic-treated LCM **(D and F)** mice were infected by oral gavage with defined mixtures of wild-type (untagged) and corresponding tagged strains. In SPF mice, both EcST131 I (*n*=6) and EcST73 (*n*=7) strains were administered at a 1:10 input ratio (tagged:untagged). In LCM (*n*=5) and antibiotic-treated LCM mice (*n*=5), EcST131 I was also administered at lower input ratios (1:1000 and 1:100000). Each data point represents one mouse, and diamonds indicate group medians. Open circles denote samples in which the tagged or untagged strain was below the limit of detection.

**Figure S19.**
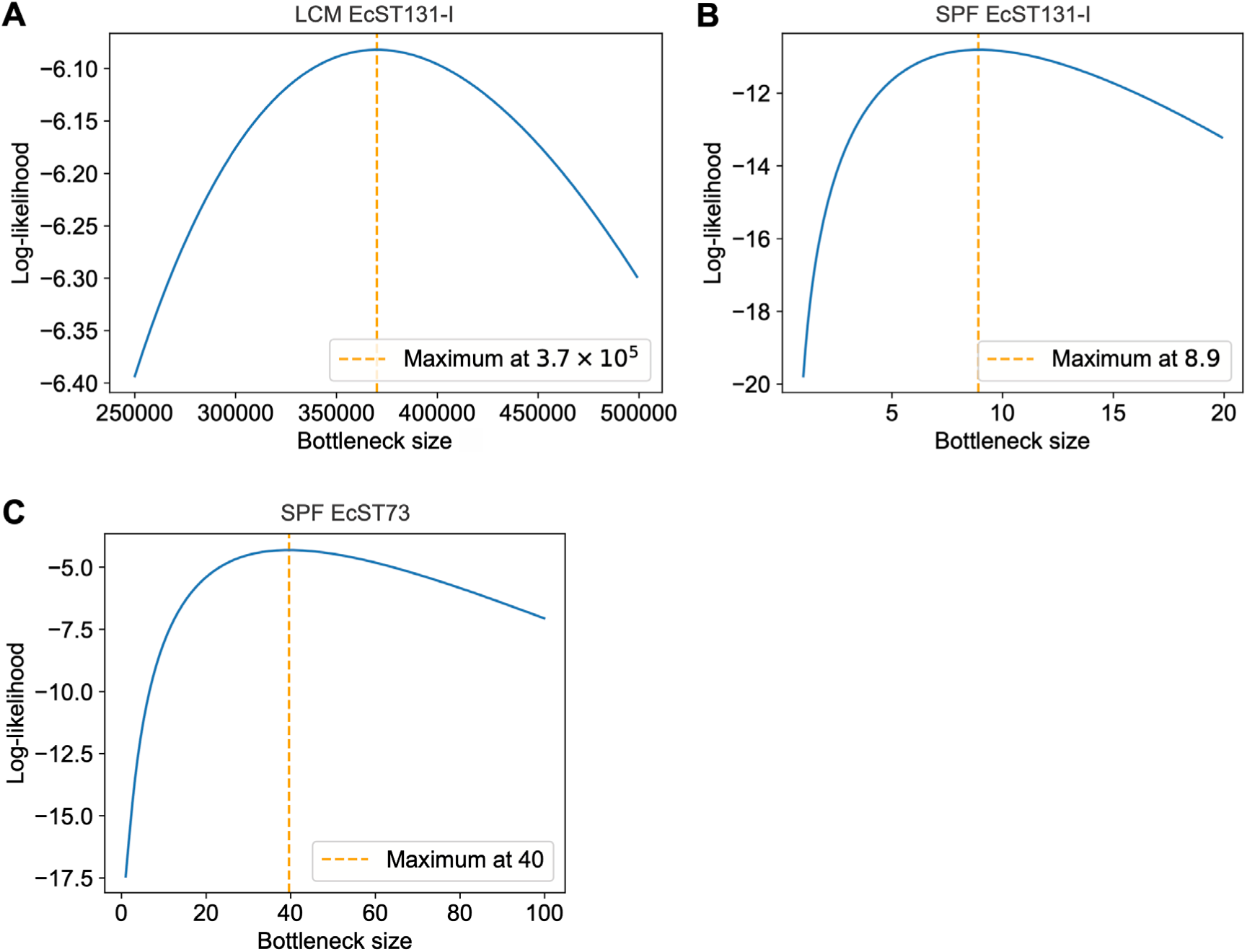
Bottleneck and extinction probability. **(A-C)** We estimate the bottleneck sizes from the experiments shown in Figure S18 using a maximum-likelihood approach described in Materials and Methods (paragraph “Bottleneck size estimation” of the section titled “Mathematical model”). **(A)** LCM mice colonized by strain EcST131 I (see **Figure S18E**); **(B)** SPF mice colonized by strain EcST131 I (see **Figure S18A**); **(C)** SPF mice colonized by strain EcST73 (see **Figure S18B**). In each case, the blue curve shows the log-likelihood versus bottleneck size, and the vertical dashed orange line indicates the bottleneck size for which this log-likelihood is maximal.

**Figure S20.**
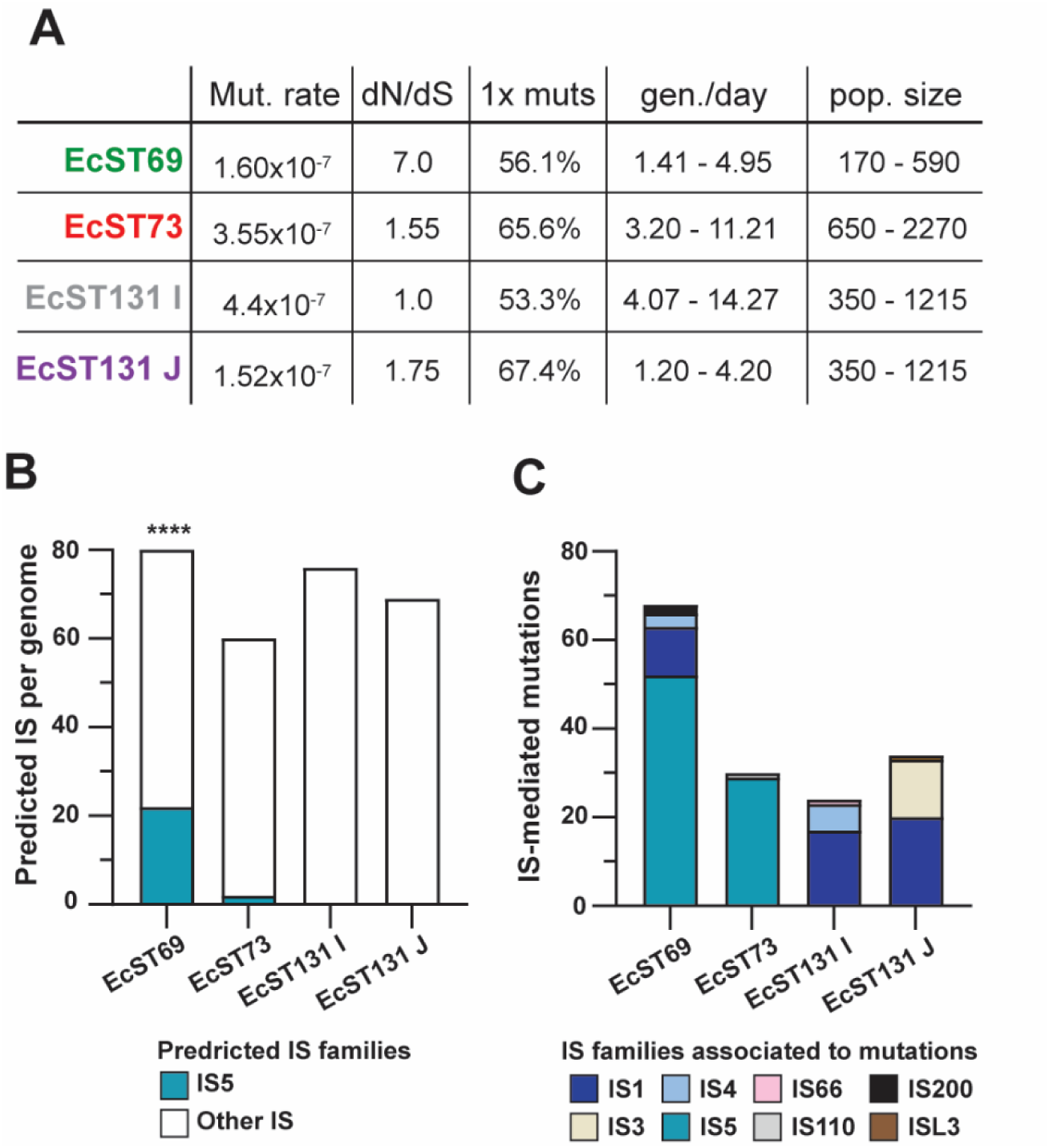
Evolution of ESBL *E. coli* strains in SPF mice shows signatures of genetic drift, IS5 transposition events, small effective population sizes, and long generation times. **(A)** Estimated mutation rates, dN/dS ratios, frequencies of one-time mutations, estimated generation time, and effective population sizes for each strain. See methods for calculation details. **(B)** EcST69 genome contains a higher number of predicted IS5 elements than the other strains, whereas the total number of predicted IS elements is similar across strains. Enrichment of predicted IS5 elements in EcST69 was assessed using Fischer’s exact test; **** *p*<0.0001. **(C)** IS5 elements account for most transposition events observed during *in vivo* evolution of EcST69. The bar plot shows the number of IS-mediated mutations (grouped by IS families) for each strain.

**Figure S21.**
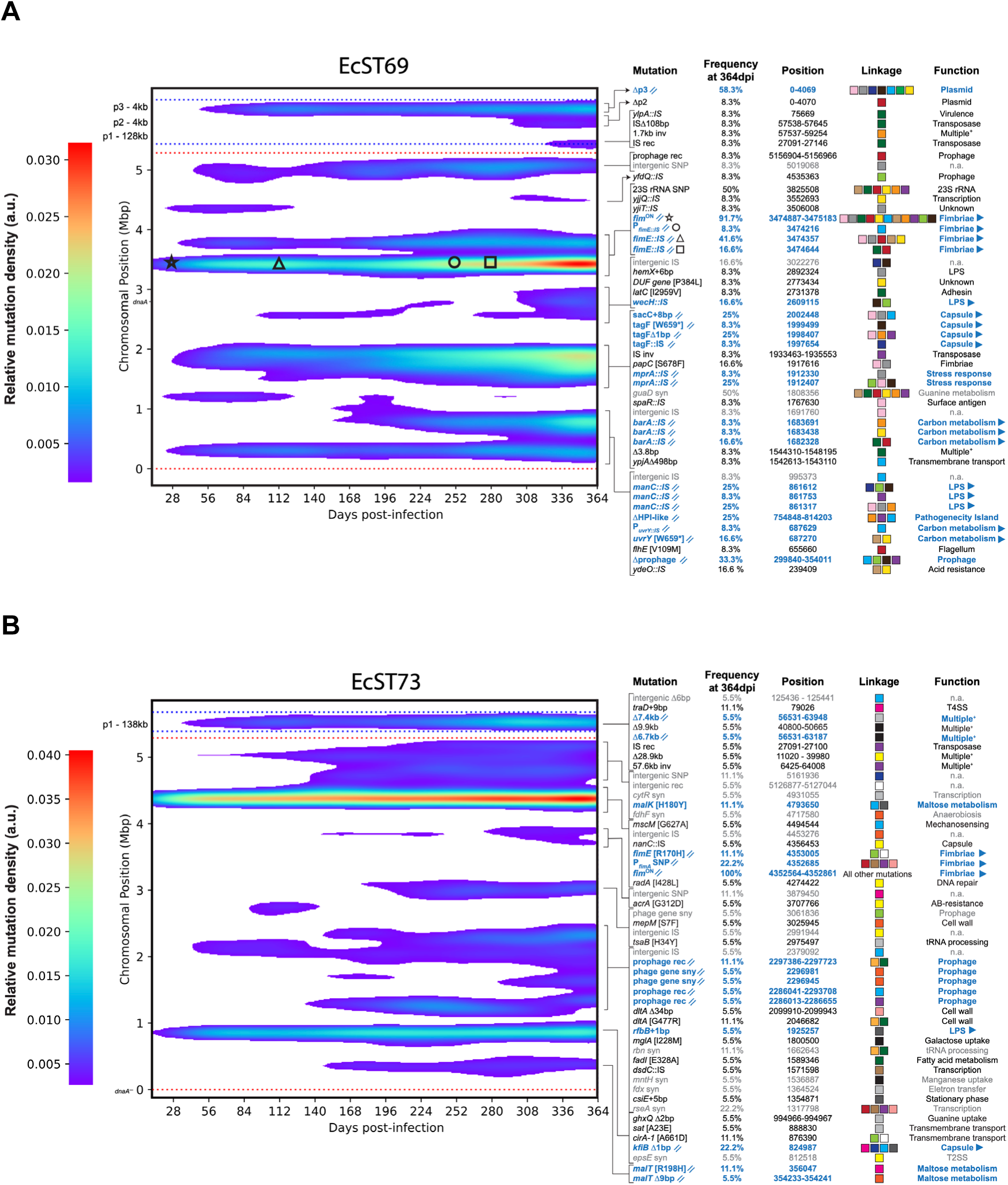
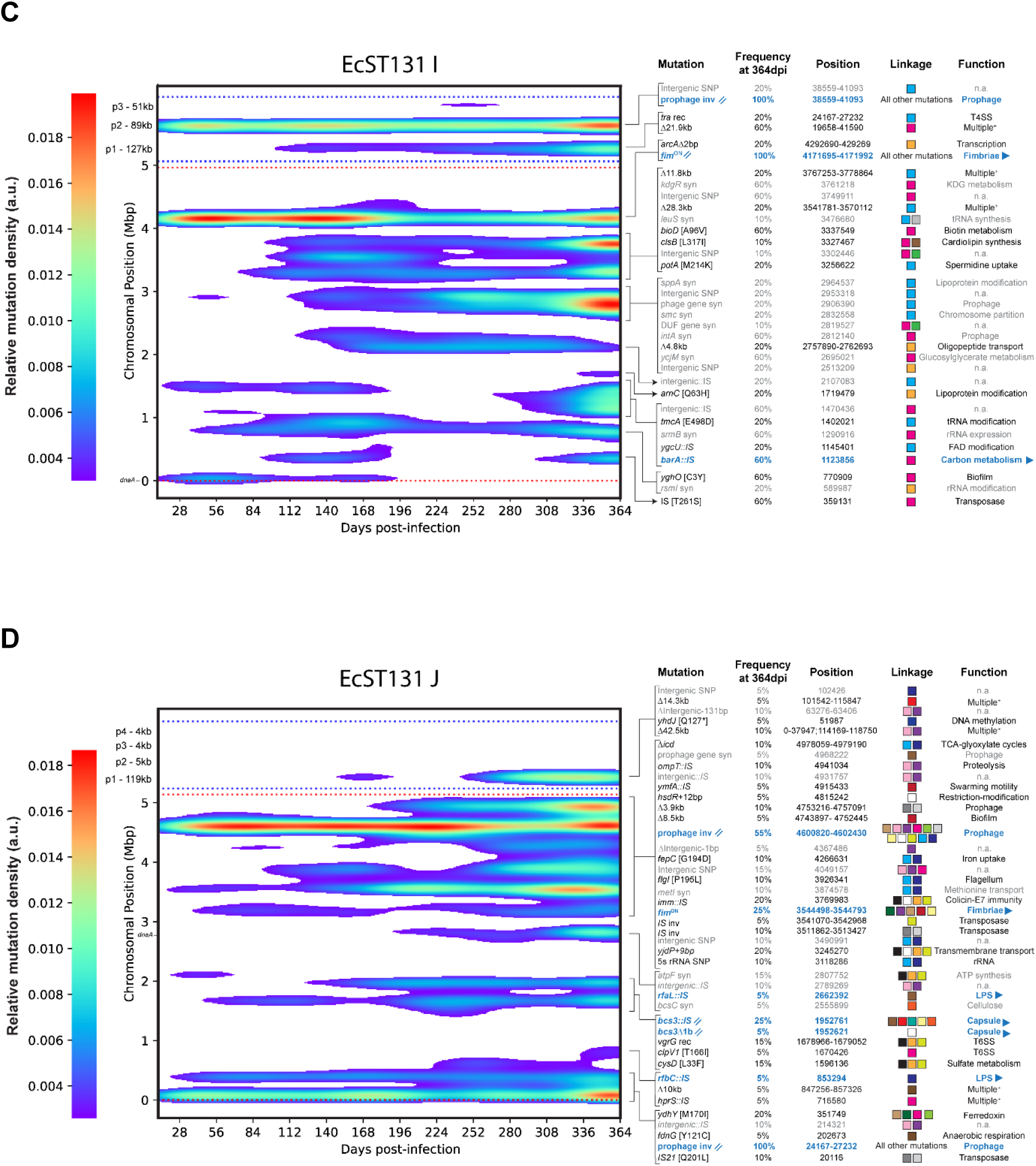
Spatiotemporal distribution of mutations during intestinal colonization of (A) EcST69, (B) EcST73, (C) EcST131 I, and (D) EcST131. **J.** Two-dimensional density maps showing the distribution of mutation occurrences over time post-infection and across genomic positions for each strain. Mutations from all clones, mice, and cages were pooled. Genomic coordinates are shown for the chromosome and plasmids, separated by spacer regions (horizontal dashed lines). Relative mutation density (arbitrary units) reflects the local concentration of mutation events after Gaussian smoothing, with higher intensity indicating regions of increased mutation clustering. // symbol denotes parallel evolution (mutations in the same genes within a strain), while ► means convergent evolution (mutations in the same genes across different strains). Blue highlights parallel and convergent evolution, while grey represents synonymous mutations or mutations in intergenic regions. Symbols marking mutations in fimbriae genes in EcST69 indicate the first appearance of the mutation. See the **supplementary data file S2** for a complete list of mutations.

**Figure S22.**
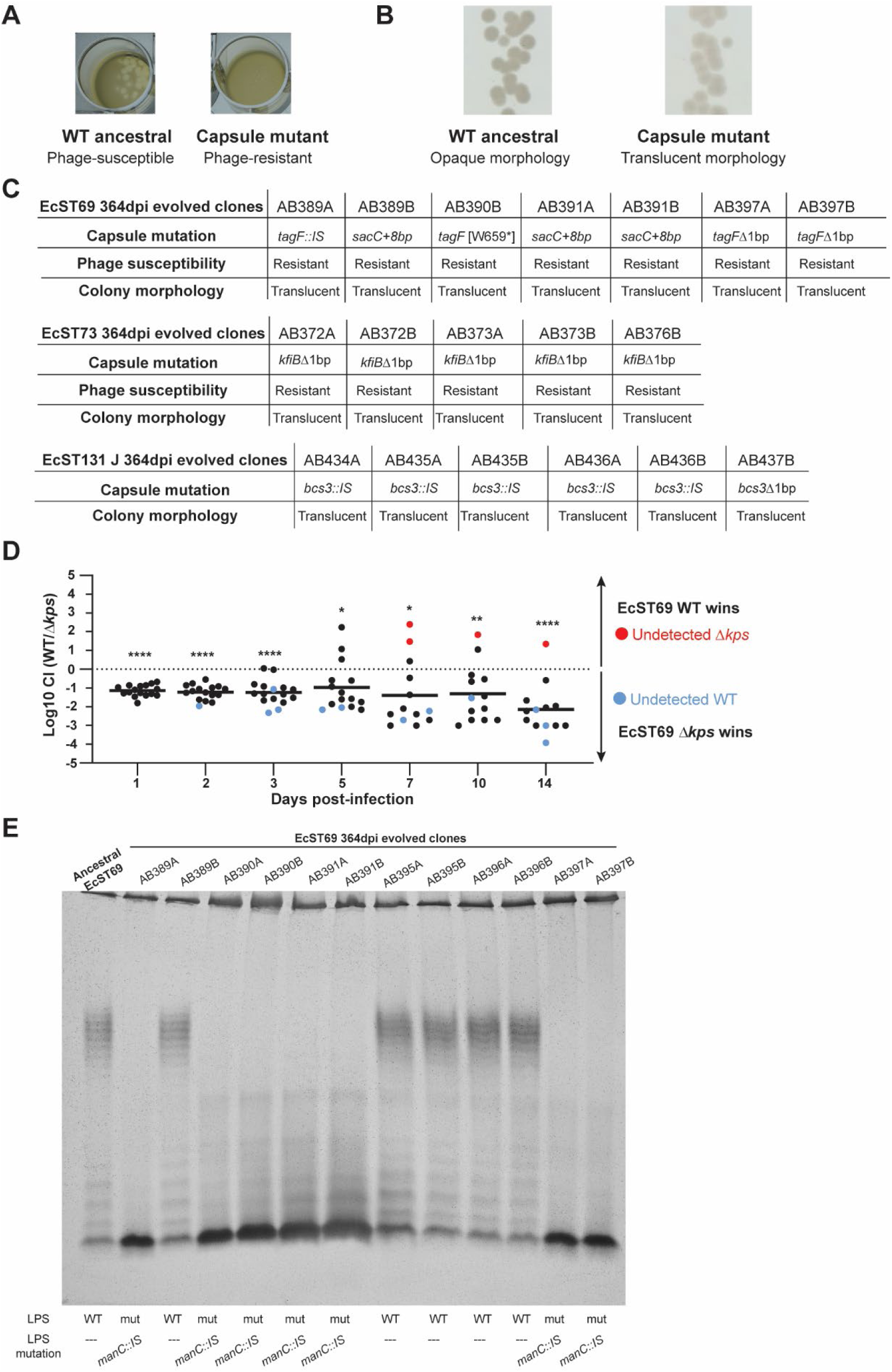
Capsule loss emerges during *in vivo* evolution of *E. coli*. **(A)** Capsule presence determines the susceptibility to phages that use the capsule as a receptor. The wild-type (WT) strain produces a capsule and is susceptible to phage infection, resulting in plaque formation in bacterial lawns, whereas a capsule-deficient mutant (Δ*kps*) is resistant and forms no plaques (representative images). **(B)** Loss of capsule results in a translucent colony morphology, in contrast to the opaque morphology of the WT (capsule-positive) strain (representative images). **(C)** All evolved clones carrying loss-of-function mutations in genes involved in the capsule biosynthesis (see the **supplementary data file S2**) displayed phage susceptibility and/or colony morphology phenotypes consistent with the capsule loss. **(D)** EcST69 Δ*kps* mutant outcompetes the WT control during gut colonization. A competition assay was performed in SPF mice over 14 days (*n*=16 mice from five independent experiments). Colored dots represent samples in which one of the strains was below the limit of detection. Mice in which both strains were undetected were excluded from statistical analyses. Statistical significance was assessed using one-sample t-test. n.s. not significant, \**p* <0.05, \*\**p* <0.01, \*\*\*\**p* <0.0001. **(E)** Mutations in genes involved in O-antigen synthesis altered lipopolysaccharide (LPS) profiles. SDS-PAGE analysis followed by silver staining revealed altered LPS banding patterns in evolved clones carrying mutations in *manC* compared with the EcST69 WT strain.

**Figure S23.**
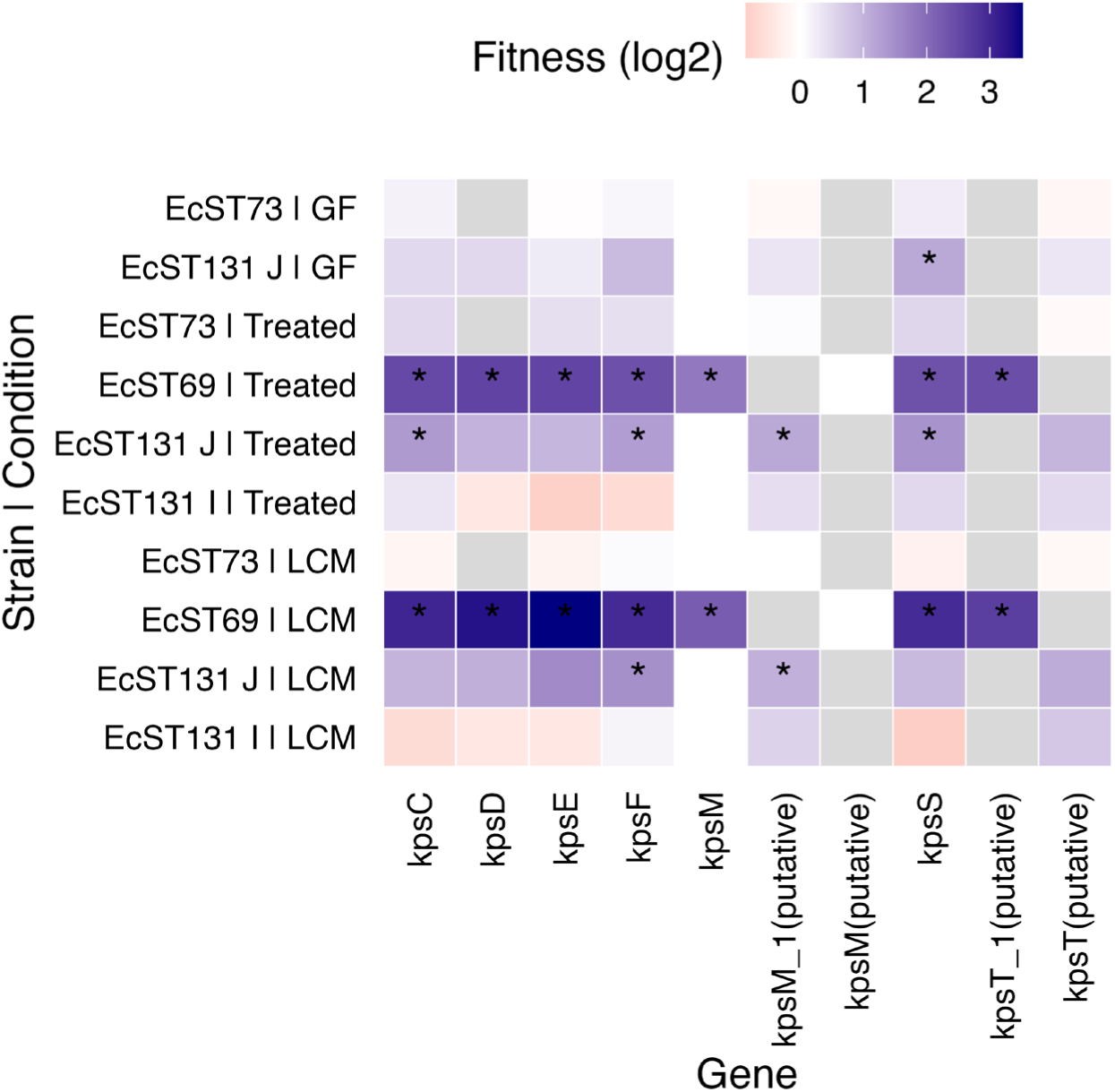
Fitness values of capsule biosynthesis genes. Heatmap shows RB-TnSeq-derived fitness scores for capsule biosynthesis genes across strains and screening conditions. Color scale indicates fold change (log_2_) in fitness, with higher values reflecting increased mutant fitness and lower values indicating fitness defects. Asterisks denote genes with statistically significant changes in fitness in the screen (see Methods).

**Figure S24.**
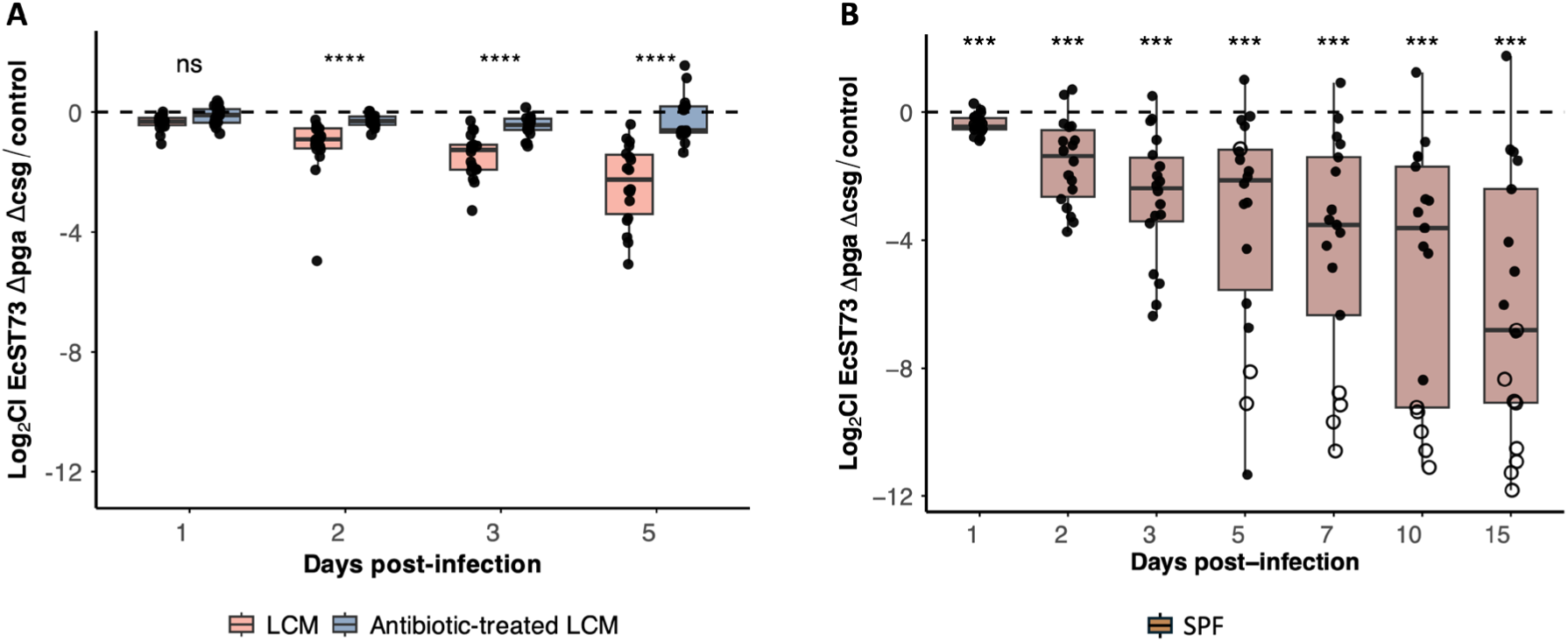
Competitive fitness of EcST73 Δ*pga*Δ*csg* mutant *in vivo* across microbiota conditions. Boxplots showing competitive indices from *in vivo* competition experiments in LCM (*n*=18), antibiotic-treated LCM (*n*=17), and conventional SPF mice (*n*=18). Each point represents an individual mouse from three independent experiments. Negative values indicate reduced fitness of the mutant relative to the control. Open circles denote mice in which the mutant was below the limit of detection. *p* values were calculated using two-sided Wilcoxon rank-sum tests **(A)** and a one-sided Wilcoxon signed-rank test **(B)**. P values were adjusted for multiple comparisons using the Holm method. ns – not significant, \**p* <0.05, \*\**p* <0.01, \*\*\**p* <0.001, \*\*\*\**p* <0.0001.

